# Age-dependent regulation of SARS-CoV-2 cell entry genes and cell death programs correlates with COVID-19 disease severity

**DOI:** 10.1101/2020.09.13.276923

**Authors:** Zintis Inde, Clarence Yapp, Gaurav N. Joshi, Johan Spetz, Cameron Fraser, Brian Deskin, Elisa Ghelfi, Chhinder Sodhi, David J. Hackam, Lester Kobzik, Ben A. Croker, Douglas Brownfield, Hongpeng Jia, Kristopher A. Sarosiek

**Affiliations:** Molecular and Integrative Physiological Sciences Program, Harvard School of Public Health, Boston, MA; John B. Little Center for Radiation Sciences, Harvard School of Public Health, Boston, MA; Harvard Program in Therapeutic Science, Harvard Medical School, Boston, MA; Image and Data Analysis Core, Harvard Medical School, Boston, MA; Integrated Cellular Imaging Core, Emory University, Atlanta, GA; Department of Surgery, Johns Hopkins University, Baltimore, MD; Division of Allergy, Immunology and Rheumatology, University of California, San Diego, CA

## Abstract

Angiotensin-converting enzyme 2 (ACE2) maintains cardiovascular and renal homeostasis but also serves as the entry receptor for the novel severe acute respiratory syndrome coronavirus (SARS-CoV-2), the causal agent of novel coronavirus disease 2019 (COVID-19)^1^. COVID-19 disease severity, while highly variable, is typically lower in pediatric patients than adults (particularly the elderly), but increased rates of hospitalizations requiring intensive care are observed in infants than in older children. The reasons for these differences are unknown. To detect potential age-based correlates of disease severity, we measured ACE2 protein expression at the single cell level in human lung tissue specimens from over 100 donors from ∼4 months to 75 years of age. We found that expression of ACE2 in distal lung epithelial cells generally increases with advancing age but exhibits extreme intra- and inter-individual heterogeneity. Notably, we also detected ACE2 expression on neonatal airway epithelial cells and within the lung parenchyma. Similar patterns were found at the transcript level: *ACE2* mRNA is expressed in the lung and trachea shortly after birth, downregulated during childhood, and again expressed at high levels in late adulthood in alveolar epithelial cells. Furthermore, we find that apoptosis, which is a natural host defense system against viral infection, is also dynamically regulated during lung maturation, resulting in periods of heightened apoptotic priming and dependence on pro-survival BCL-2 family proteins including MCL-1. Infection of human lung cells with SARS-CoV-2 triggers an unfolded protein stress response and upregulation of the endogenous MCL-1 inhibitor Noxa; in juveniles, MCL-1 inhibition is sufficient to trigger apoptosis in lung epithelial cells – this may limit virion production and inflammatory signaling. Overall, we identify strong and distinct correlates of COVID-19 disease severity across lifespan and advance our understanding of the regulation of ACE2 and cell death programs in the mammalian lung. Furthermore, our work provides the framework for potential translation of apoptosis modulating drugs as novel treatments for COVID-19.

## INTRODUCTION

The novel coronavirus disease (COVID-19) pandemic has caused infection of over 25.6 million individuals globally as of 31 August 2020^2^. Disease severity is typically lower in pediatric patients than adults (particularly the elderly), but higher rates of hospitalizations requiring intensive care are observed in infants than in older children^3–5^. The causal agent for COVID-19, the novel severe acute respiratory syndrome coronavirus (SARS-CoV-2), infects host cells through interaction with the cell surface proteins angiotensin-converting enzyme 2 (ACE2) and the transmembrane serine protease TMPRSS2^1^. For SARS-CoV-2 infection to occur, the spike protein of a viral particle must bind ACE2 and undergo cleavage by TMPRSS2 to allow fusion with the host plasma membrane and cell entry^1^. In the lung, where this process is known to occur, developmental processes in early life require coordinated regulation of gene expression to ensure proper formation and maturation of airways and alveoli^6^. It remains unknown how these developmental processes affect ACE2 expression and potentially contribute to differences in COVID-19 disease severity across lifespan.

In addition to its role in SARS-CoV-2 infection, ACE2 regulates vascular homeostasis as part of the renin-angiotensin system^7^. In this normal physiological role, ACE2 is expressed on cell membranes in a number of tissues outside the lung. Several single-cell transcriptomic analyses have been performed to determine the tissue and cell type distribution of *ACE2* expression^8–11^. Although they have identified cell types that may be susceptible to infection, these studies generally lack data across different ages and are unable to examine ACE2 translation and localization. Protein-level measurements, therefore, are required to determine whether *ACE2* mRNA expression correlates robustly with translated, membrane localized protein expression.

Viral entry represents just one of several steps in COVID-19 pathogenesis, each of which involves genes that may potentially be developmentally regulated in the lung and elsewhere. Apoptotic and non-apoptotic host cell death pathways also play important roles as they can modulate disease pathogenesis after viral infection^12,13^. Specifically, coronavirus-infected cells typically experience endoplasmic reticulum (ER) stress due the excessive production of infectious virions and consequently activate the unfolded protein response (UPR) to adapt to this stress^14^. If the ER stress is overwhelming, the cell undergoes intrinsic apoptosis (programmed cell death) by upregulating pro-apoptotic proteins – this serves as a host defense mechanism by arresting virion production. Interestingly, virally-encoded proteins may suppress this process during infection to enhance virion production^15–17^. Apoptosis is controlled by the members of the BCL-2 family of proteins, which have pro-death or pro-survival roles in modulating mitochondrial release of cytochrome c, the commitment point for intrinsic apoptosis^18^. We have previously found that apoptosis is dynamically regulated during lifespan in multiple organs as they grow and mature^19^ -this could impact the degree to which cells tolerate ER stress and thus extent of virion production upon infection.

Herein, we investigate how expression of viral entry and cell death genes in the lung might vary among different age groups and how it relates to known differences in disease severity. We find age-based differences in ACE2 expression in mouse lung tissue that correlate with varying severity of human COVID-19 symptoms in adults, children, and neonates. Importantly, we validate these differences by measuring expression of ACE2 at the protein level across over 100 normal lung tissue specimens, extending the results of previous transcriptomic analyses. Notably, we detected increased expression of this viral receptor on multiple cell types in the lung during key life stages and extreme levels of inter- and intra-individual heterogeneity that may influence disease course across populations. Finally, we find that apoptosis is dynamically regulated in the postnatal lung to create periods of heightened apoptotic priming and dependence on pro-survival proteins that can limit the extent of virion production upon SARS-CoV-2 infection and has implications for clinical translation.

## RESULTS

In mouse and human lung, developmental processes in postnatal life require coordinated regulation of gene expression in epithelial and endothelial cells to ensure proper formation and maturation of airways and highly vascularized alveoli^6^. The importance of ACE2 in maintaining vascular homeostasis suggests that it may also be dynamically regulated in the lung throughout life, which may in turn impact cellular entry and infection by coronaviruses including SARS-CoV-2^20^. Although previous reports have demonstrated expression of *ACE2* in various adult cell types within the lung as well as in small intestine, kidney, and testis tissue^8–11^, we sought to characterize how ACE2 expression changes at the transcript and protein levels across the mammalian lifespan.

To detect potential changes in expression, we stained for ACE2 and cell markers of interest in two tissue microarrays (TMA) comprising over 100 normal lung specimens from individuals ranging from 9 to 75 years of age. Due to the pronounced autofluorescence of red blood cells (Figure 1A), we identified and segmented cells that stained positive for DAPI to evaluate total cellular ACE2 expression levels in 1,408,152 nucleated cells (Figure 1B). We first compared ACE2 expression in all lung cells across all individuals (Figure 1C) and found that the percentage of ACE2-expressing cells in the lung increased with age but was profoundly heterogeneous (Figure 1D-F). In fact, while some normal lung tissues from similarly aged individuals contained an extensive number of ACE2^+^ cells (>70% of DAPI^+^), others expressed very few cells in the distal lung that can be infected by SARS-CoV-2 (<5% of DAPI^+^). We attempted to identify the cell types in which ACE2 was expressed by co-staining for podoplanin (PDPN), a marker commonly expressed on alveolar type 1 (AT1) cells that are responsible for gas exchange. Co-staining with PDPN showed increasing ACE2 expression in PDPN^+^ cells with age and again demonstrated extensive heterogeneity within and between individuals (Figure 1G-J). However, while we observed ACE2^+^ PDPN^+^ cells, their morphology was cuboidal instead of thin and elongated as would be typically found in AT1 cells. This suggests that they may in fact be alveolar type 2 (AT2) cells, which secrete surfactant and regenerate the alveolar epithelium, including AT1 cells, after injury^21^. Also, while PDPN was detected in these cells, it was observed primarily in intracellular puncta, rather than on the luminal surface found commonly on AT1 cells. These unexpected differences in PDPN staining may be due to the TMA tissue being sourced from patients at autopsy or from normal lung tissue adjacent to lung tumors (see methods); either of these, as well as environmental or disease-associated factors, could potentially reduce the specificity of PDPN staining for AT1 cells or alter AT1 cell morphology^22,23^. Finally, although lung specimens from male donors tended to have higher expression of ACE2 in nucleated (Figure S1J) and PDPN^+^ (Figure S1K) cells than female donors, the differences were not statistically significant and similar age-based trends were evident for both sexes.

**Figure 1:**
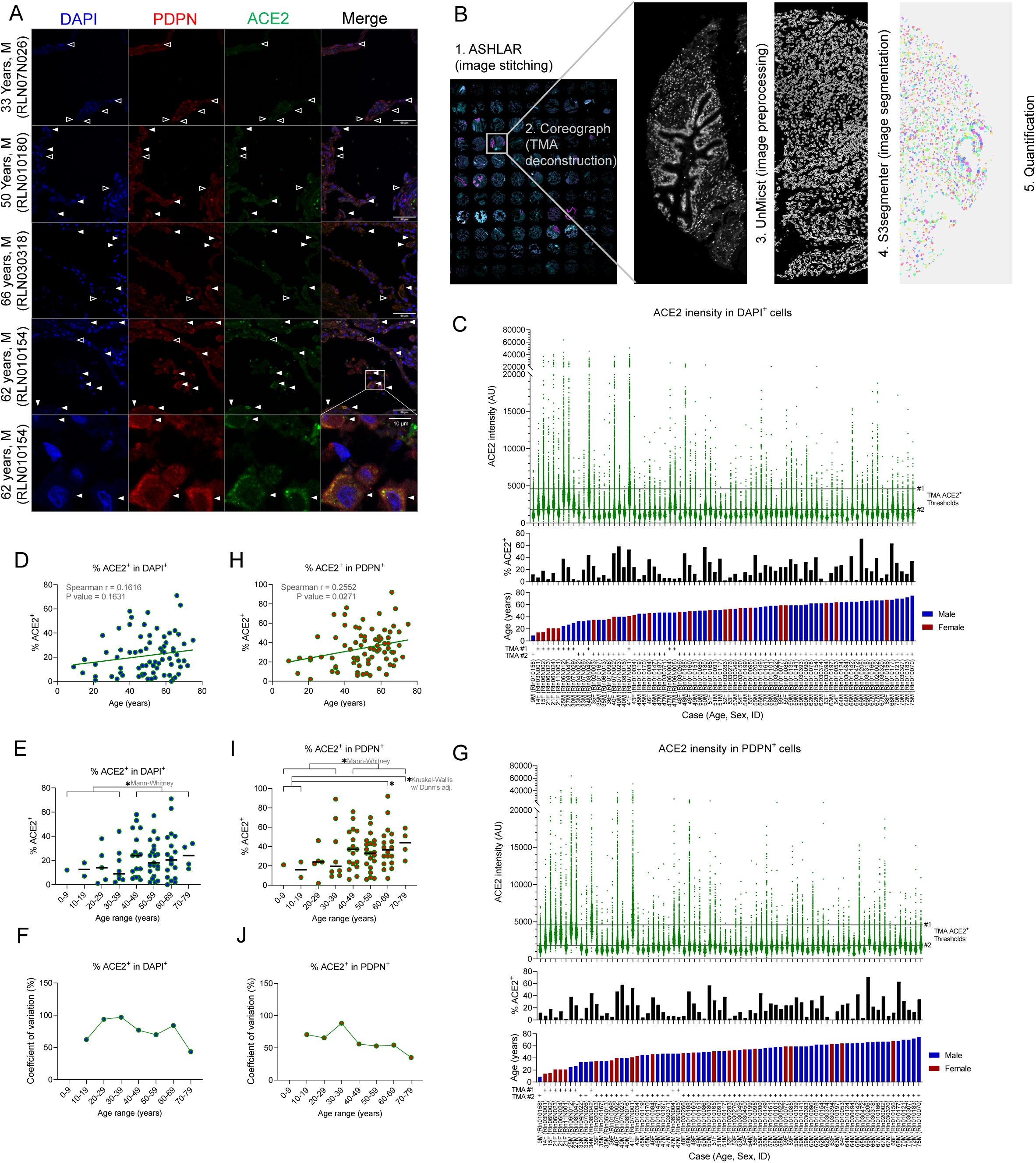
ACE2 expression in all nucleated and PDPN^+^ cells. (A) Images of immunofluorescence staining for ACE2 and PDPN in lung tissue from humans at the indicated ages. Filled arrowheads indicate examples of cells that are positive for indicated marker (PDPN) and also positive for ACE2 while empty arrowheads are positive solely for the indicated marker but not ACE2. Scale bar = 50 µm, or 10 µm for magnified region of interest. (B) Schematic of image processing and quantification pipeline for human tissue microarrays (TMAs). Images are first acquired as single tiles on a widefield fluorescence microscope and are stitched (ASHLAR). The TMA is separated into single organized cores (Coreograph) and the DAPI channel of each core is preprocessed to accentuate nuclei boundaries for easier segmentation (UnMicst). Based on this intermediate image, the nuclei and corresponding cytoplasmic regions are compartmentalized as separate indexed objects (S3segmenter) and the mean/median intensity of each is quantified. (C) ACE2 intensity measurements in DAPI^+^ cells within lung cores (top) with the percentage of cells being positive (>2250 AU intensity) for ACE2 (middle) and the age and sex of patients (bottom). (D, E) Correlation analysis comparing age with percent ACE2 positivity in all cells in normal human lung tissue. (F) Coefficient of variation for ACE2^+^ percentages in TMA cores from the age ranges shown in D. (G) ACE2 intensity measurements in PDPN^+^ cells within lung cores (top) with the percentage of cells being positive (>2250 AU intensity) for ACE2 (middle) and the age and sex of patients (bottom). (H, I) Correlation analysis comparing age with percent ACE2 positivity in PDPN^+^ cells in normal human lung tissue. (J) Coefficient of variation for ACE2^+^ percentages in TMA cores from the age ranges shown in H.

We next assessed levels of ACE2 expression specifically in AT2 cells by staining for aquaporin 3 (AQP3) in the two TMAs. In contrast with PDPN^+^ cells, expression of ACE2 did not increase with age in AT2 cells in these specimens (Figure 2A-D), although several of the specimens were removed from analysis due to inadequate specimen quantity or quality (see methods). The expression of ACE2 was again highly heterogeneous (Figure 2E) with similarly aged individuals expressing a wide range of ACE2 positivity (from ∼0 to >90%). Since our TMAs did not contain specimens from neonates or infants, we next sought to obtain neonatal lung tissue to assess the early-life expression of ACE2 in human lung tissue. We detected ACE2 expression on the apical surfaces of airway epithelial cells (likely club cells) in infant (age 3-5 months) lung tissue (Figure 2F, G). We also detected ACE2^+^ cells outside of the airway, where it seemed to be expressed on epithelial cells.

**Figure 2:**
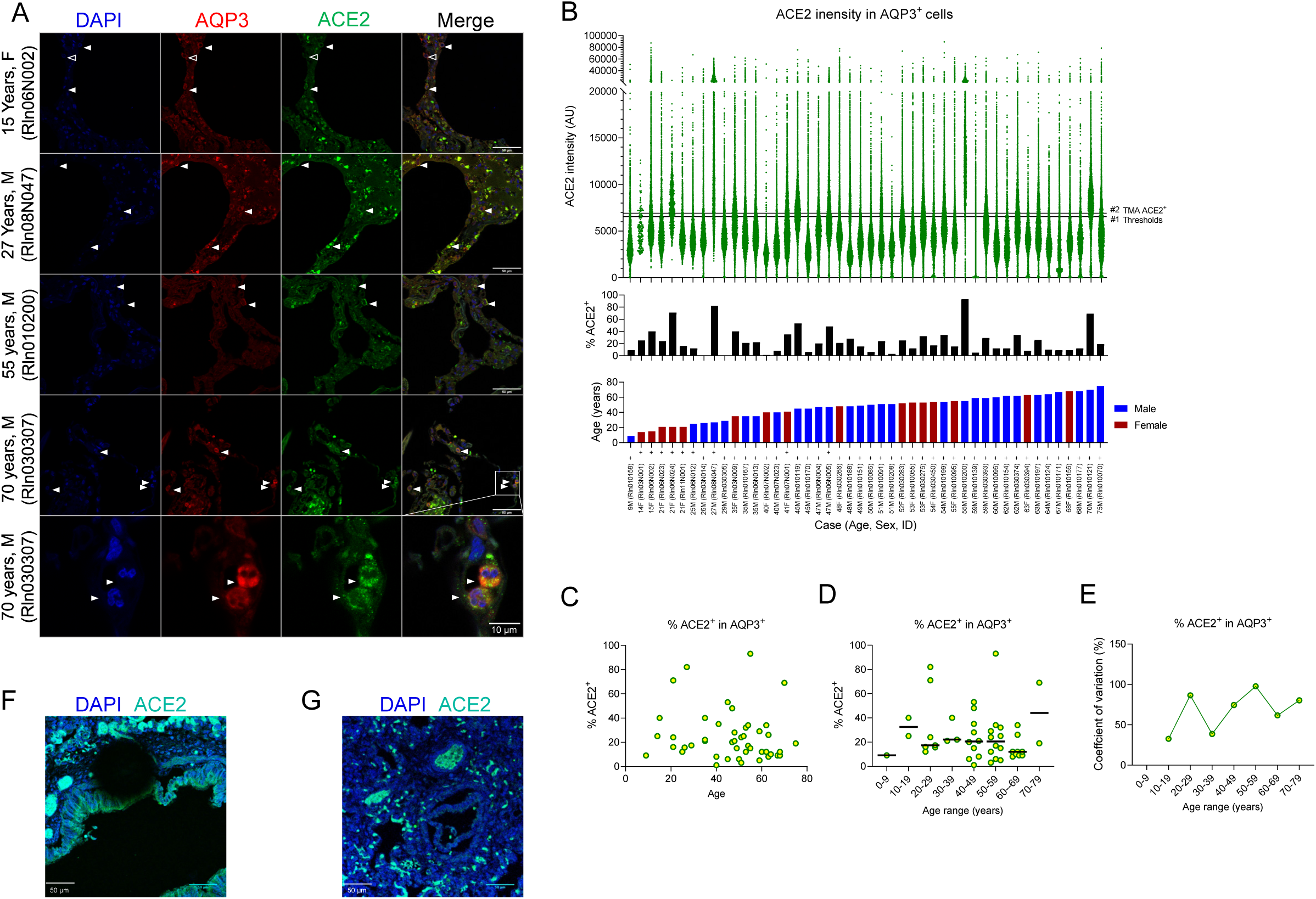
ACE2 expression in human alveolar type II cells and neonatal lung. (A) Images of immunofluorescence staining for ACE2 and AQP3 in lung tissue from humans at the indicated ages. Filled arrowheads indicate examples of cells that are positive for indicated marker (AQP3) and also positive for ACE2 while empty arrowheads are positive solely for the indicated marker but not ACE2. Scale bar = 50 µm, or 10 µm for magnified region of interest. (B) ACE2 intensity measurements in AQP3^+^ cells within lung cores (top) with the percentage of cells being positive for ACE2 (middle) and the age and sex of patients (bottom). (C, D) Correlation analysis comparing age with percent ACE2 positivity in AT2 cells in normal human lung tissue. (E) Coefficient of variation for ACE2^+^ percentages in TMA cores from the age ranges shown in D. (F, G) Images of immunofluorescence staining for ACE2 in neonatal human lung tissue (age 3-5 months). Scale bar = 50 µm.

Given the heterogeneity we observed in human ACE2 expression, we reasoned that examining ACE2 expression in inbred mouse strains might minimize inter-individual variation and validate the age-dependent changes we observed in human tissue. Although some temporal and spatial aspects of lung maturation vary between the mouse and human, the stages of lung development are similar across all mammalian species^24^. In both the human and murine lung, alveologenesis (formation of highly vascularized alveoli, which are the primary gas-exchange units in the lung), begins prenatally but isn’t complete until young adulthood^24^ (∼P36 in mice, ∼21 years of age in human^25,26^). Furthermore, mouse models have become critical for research into coronavirus infection pathogenesis. In particular, these models have shown that host age, virus-host protein-protein interactions, and expression levels of viral receptors can all affect the “effective dose” of viral particles experienced by cells in the lung; this dosage can in turn determine the severity of symptoms (Table 1)^27^. Therefore, we queried expression of both *Ace2* and the serine protease *Tmprss2* (a host cell protease required for SARS-CoV-2 cell entry after binding to ACE2^20^) across 156 samples comprising 31 microarray experiments using a tool for combined analysis of published microarray data^28^. We found that lung tissue collected from newborn mice (postnatal day 0-3 [P0-P3]) exhibited higher *Ace2* expression levels than P4-15 adolescent mice and that expression increased steadily after adolescence through to advanced age (P256+) (Figure 3A). A similar expression pattern was observed for *Tmprss2*, although the magnitude of differences was smaller (Figure S2A). We confirmed these results in a separate set of 66 samples from 5 additional experiments using a distinct microarray platform (Figure S2B-C).

**Table 1:**
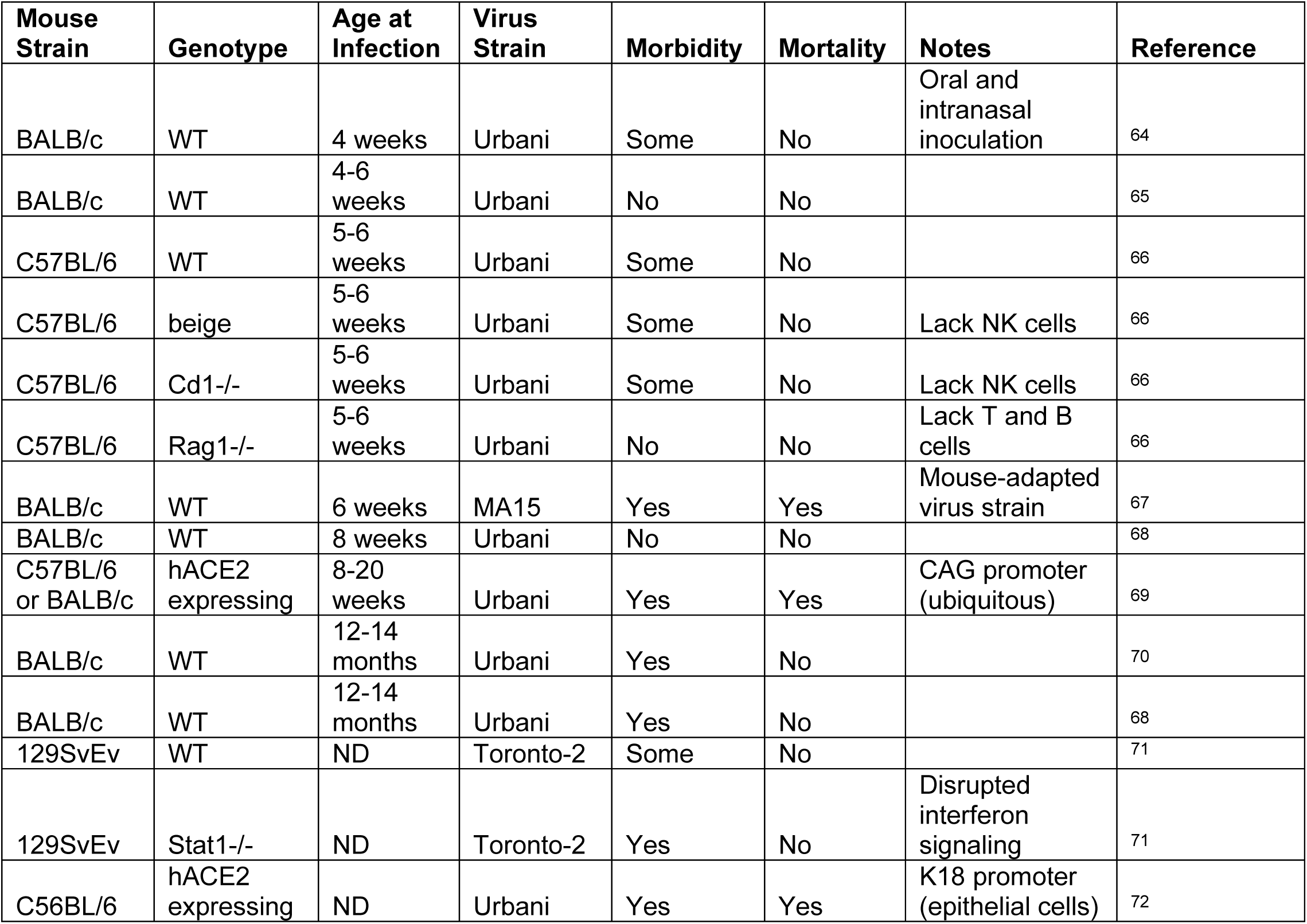
Selected mouse models of SARS-CoV infection show effect of age and *ACE2* expression

**Figure 3:**
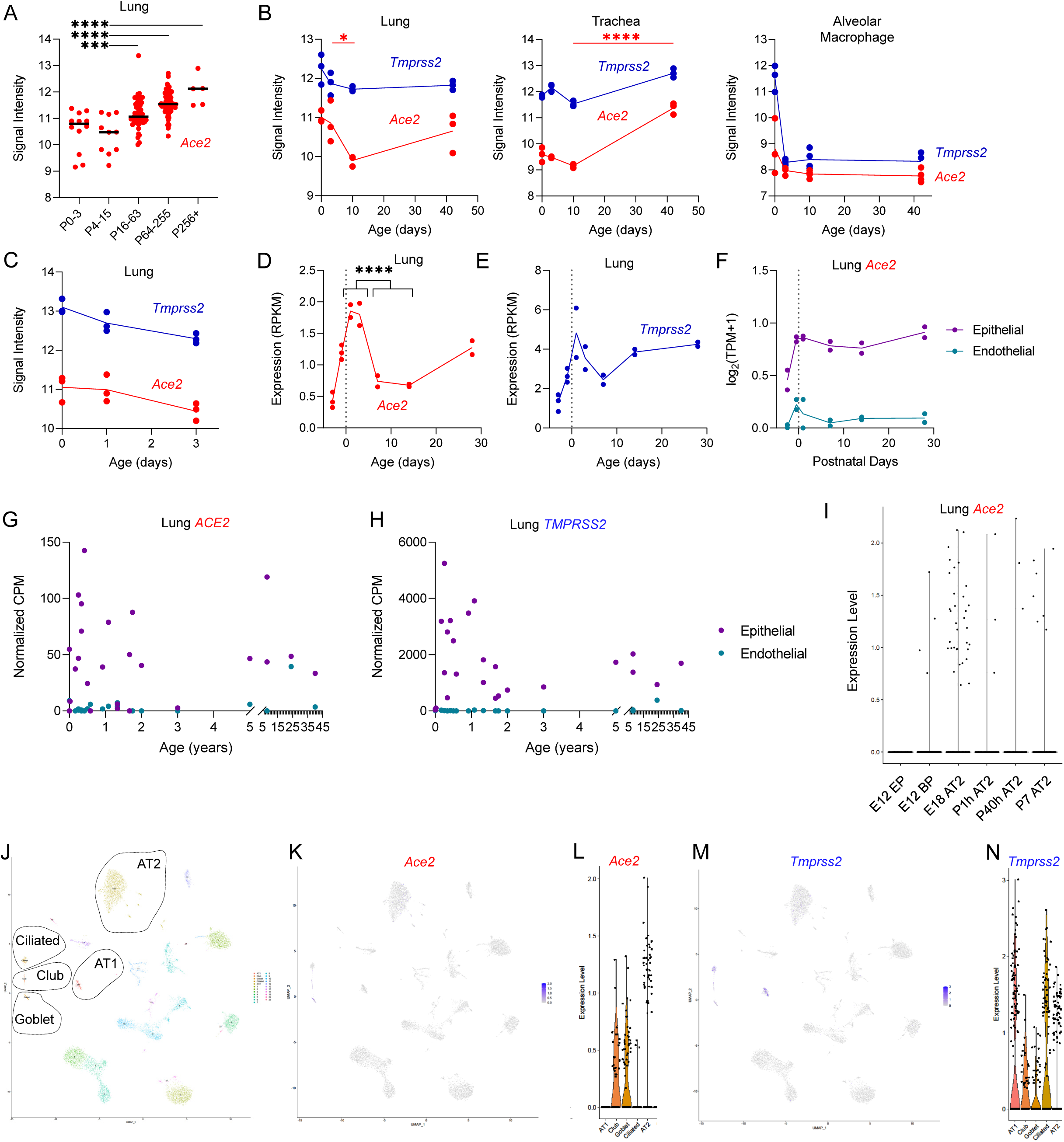
ACE2 and TMPRSS2 are dynamically regulated postnatally in the mouse and human lung. (A-C) Mouse microarray gene expression data. (A) Collected data from experiments conducted on mouse lung tissue of the indicated age ranges using the Affymetrix Mouse Genome 430 2.0 array. Comparison between groups was conducted by one-way ANOVA with Holm-Sidak’s adjustment. (*, p<.05; ***, p<.001; ****, p<.0001) (B) Expression of *Ace2* and *Tmprss2* in mouse lung, trachea, and macrophages collected by bronchoalveolar lavage (C) Expression of *Ace2* and *Tmprss2* in whole lung. (D, E) RNA sequencing data showing *Ace2* (D) and *Tmprss2* (E) expression in whole lung. (F) RNA sequencing data showing *Ace2* expression in FACS-sorted cell types from mouse lung. (G, H) RNA sequencing data showing *ACE2* (G) and *TMPRSS2* (H) expression in FACS-sorted cell types from human lung. (I) Single cell RNA sequencing showing *Ace2* expression in AT2 cells and their precursors (EP: epithelial progenitor; BP: bipotent progenitor) in mouse lung at the indicated pre- and postnatal timepoints. (J, K, M) UMAP plots of adult mouse lung scRNA-seq, showing cell type clusters (J) as well as expression of Ace2 (K) and *Tmprss2* (M). (L, N) Dot plot for selected cell types are shown alongside corresponding UMAP plots.

For individual experiments from this collected dataset that included detailed time course data, patterns matched those observed in the combined data. Analysis of a study of gene expression in P0, P3, P10, and P42 mice showed only minor changes in *Tmprss2* levels in whole lung and trachea, but *Ace2* expression was lower at P10 than all other timepoints in both tissues^29^ (Figure 3B). *Ace2* and *Tmprss2* were expressed at high levels in alveolar macrophages at P0 before declining substantially at P3 and later. Similarly, *Tmprss2* and *Ace2* levels were highest immediately after birth in a study of lung tissue at P0, P1, and P3^30^ (Figure 3C). Another study of tissue from P1, P8, and P28 mice did not show a decline in *Ace2* levels at P8, but did show elevated expression at P28^31^ (Figure S2D). Thus, across a diverse set of microarray datasets, lung *Ace2* levels were relatively high immediately after birth, significantly lower during adolescence, and increased in adulthood, reaching their peak at advanced age.

To complement these microarray data, we queried the Lung Gene Expression Analysis (LGEA) database of mouse RNA-seq data across developmental timepoints^32^. We again found that expression of *Ace2* and *Tmprss2* peaked around birth, declined in young mice, and increased again in older mice (Figure 3D, E). In sorted cells from the LGEA dataset, *Ace2* levels in epithelial cells were high at birth and were reduced shortly after, followed by an upregulation when alveologenesis is nearing completion^24^ (Figure 3F). *Ace2* expression in vascular endothelial cells peaked at birth and then decreased by P30. *Tmprss2* was similarly expressed during this time period (Figure S2E).

In our human lung TMA experiments, we had examined ACE2 expression at the protein level but did not analyze *ACE2* mRNA transcripts. To compare with mouse transcriptional data, we analyzed the LungMAP database, which contains RNA-seq data for sorted human lung cell types from donors of varying age^33^. Consistent with murine data, human epithelial cell expression of *ACE2* and *TMPRSS2* was detected in neonates, infants, children, and young adults (24-40 years of age), with the highest levels evident in infant lung tissue (Figure 3G, H). The heterogeneity between individuals was consistent with the protein-level heterogeneity observed in our analysis of the TMAs. Endothelial cells within the human lung expressed higher *ACE2* in adults than at other ages, although the sample size was limited.

These mouse and human data showed similar trends in expression by bulk RNA-seq, but the sorting strategy employed in this analysis combined epithelial cells of various subtypes. Previous reports have demonstrated expression of *Ace2* specifically in AT2 epithelial cells^8,10^; we sought to determine whether this expression might vary with time. In single cell RNA-seq data for AT2 cells and their embryonic precursors, we found *Ace2* and *Tmprss2* can be expressed in AT2 cells as early as embryonic day 12 through to P7 (Figure 3I and S2F). In middle adulthood (4 months), single-cell RNA-seq data showed that AT2 cells may express *Ace2* and *Tmprss2*, but broader expression was detected in bronchiolar epithelial cells including club and goblet cells (Figure 3J-N). These data strongly suggested that *Ace2* is dynamically expressed in major lung cell types during postnatal life, potentially modulating (in concert with *Tmprss2* and other proteases) the number and type of cells susceptible to SARS-CoV-2 infection (summarized in Table 2).

**Table 2:** Age-dependent regulation of ACE2, TMPRSS2 and apoptosis in mouse lung.

| Mouse age: |  | P0-P3<br>(Neonate) | P7-P15<br>(Adolescent) | P28-P100<br>(Young Adult) | P250+<br>(Late Adult) |
| --- | --- | --- | --- | --- | --- |
| Ace2 Lung |  | Moderate | Low | Moderate | High |
| ACE2 in Lung<br>Cells | All (DAPI+) | Moderate | Low | Moderate | Very High |
|  | SCGB1A1+ | Moderate | Moderate | High | Very High |
|  | PDPN+ | Low | Low | Low | Moderate |
|  | AQP3+ | Low | Low | Low | Very High |
| Tmprss2 Lung |  | Moderate | Low | Moderate | High |
| Apoptotic Priming |  | High | High | Moderate | Moderate |
| BCL-2 Family<br>Gene Expression | Pore-Forming | Low | Moderate | Moderate |  |
|  | Pro-Apoptotic | High | Moderate | Low |  |
|  | Pro-Survival | High | Moderate | Moderate |  |

For a more direct comparison with our TMA staining, we next sought to investigate ACE2 protein expression dynamics directly in mouse lung tissue across lifespan. We stained for ACE2 and relevant cell markers in lung tissue from newborn (P0), young (P7), adult (3 months) and late adult (11 months) mice (Figure 4). We detected slightly elevated levels of ACE2 across all nucleated cells immediately after birth, which were lower at P7 and increased by adulthood (3 months) (Figure 4A-B). We found that the higher perinatal ACE2 levels were at least in part attributable to its expression in AQP3 positive AT2 cells (Figure 4A, C), which was consistent with our gene expression analysis. Unexpectedly, we also found that ACE2 protein expression was strongly increased by late adulthood in AT2 cells, and that it was expressed throughout the cell including the nucleus (Figure 4A). We also stained for secretoglobin 1A1 (SCGB1A1), a marker for club cells within the airway (Figure 4D, E), and found that expression levels were high at P0, downregulated by P7, and increased at later ages. Finally, in agreement with our human TMA data, faintly ACE2^+^ cells were observed that could be AT1 given their proximity to PDPN (Figure 4, G). However, an AT2 co-stain as well as another AT1 marker (e.g. receptor for advanced glycation end products – RAGE) would be required to confirm whether they are *bona fide* AT1 cells. In summary, we detected age-based differences in ACE2 expression in multiple cell types within the lung (Figure 4H), which may delineate the pool of cells that can be infected by SARS-CoV-2 at different ages. Intriguingly, changes in ACE2 expression, which are heterogeneous but broadly consistent across mouse and human samples, are correlated with COVID-19 disease severity in human patients (Figure S1A-C).

**Figure 4:**
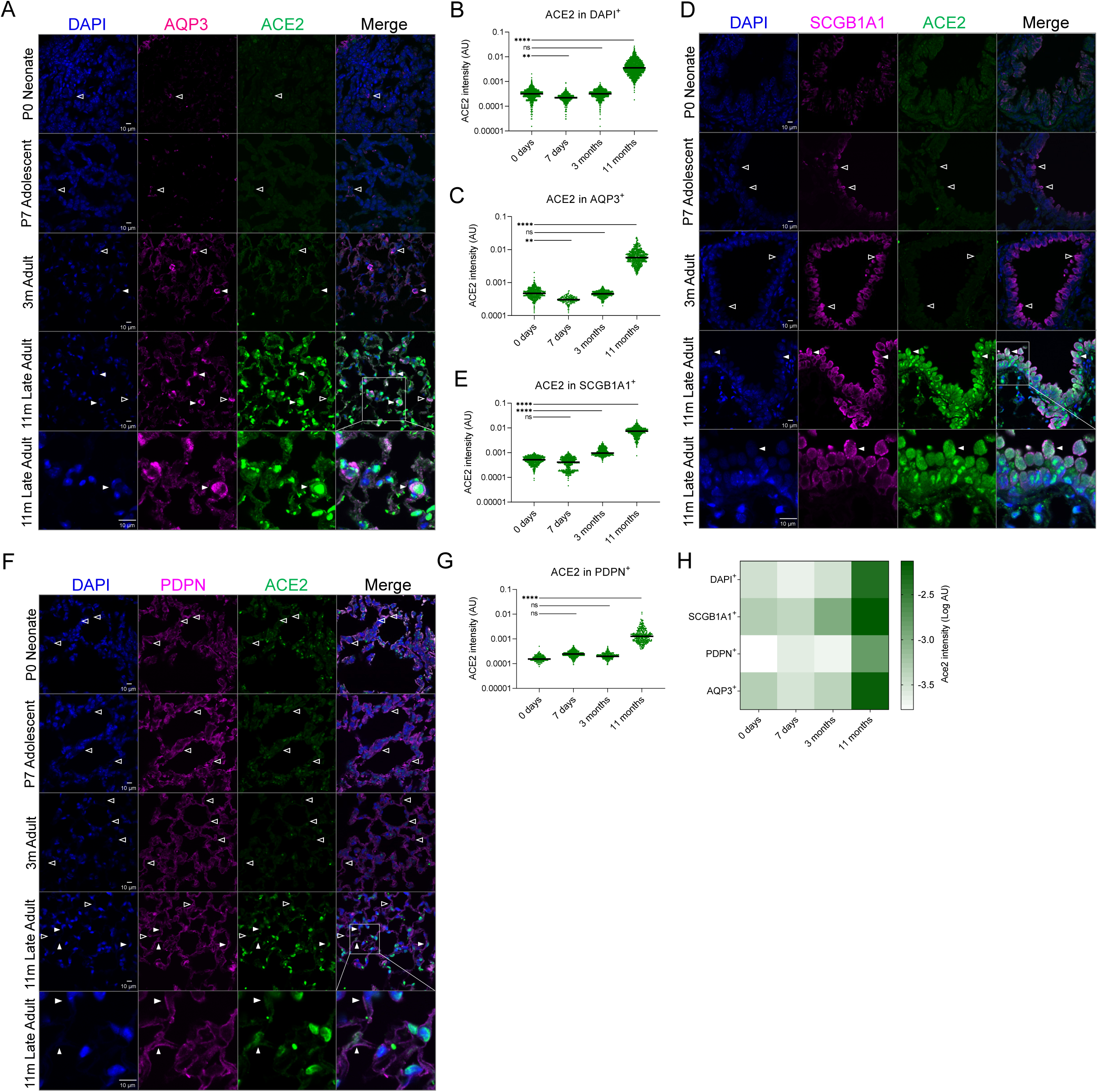
Dynamic ACE2 expression in mouse postnatal lung. (A, D, F, H) Images of immunofluorescence staining for ACE2 and other indicated proteins in lung tissue from mice at the indicated ages. Filled arrowheads indicate examples of cells that are positive for indicated marker and also positive for ACE2 while empty arrowheads are positive solely for the indicated marker but not ACE2. Scale bar = 10 µm. (B, C, E, G, I) Quantification of ACE2 fluorescence in all cells (B) or in cells with indicated marker (C, E, G, I). Comparison between groups was conducted by one-way ANOVA with Holm-Sidak’s adjustment (**, p<.01;***, p<.001; ****, p<.0001). (J) Comparison of average ACE2 staining intensities across cell types tested.

These patterns suggested that higher lung ACE2 expression in infancy and late adulthood would broaden the pool of cells that can be potentially infected by SARS-CoV-2 at those ages. However, in response to cellular stress induced by viral infection, cells frequently trigger apoptosis as a host defense mechanism^12,13^. In the case of coronaviruses, active infection and stimulation of virion production typically results in proteotoxic stress and consequent activation of the unfolded protein response (UPR)^16^, which is also a potent inducer of apoptotic cell death^34,35^. Apoptosis can be suppressed during infection by virally-encoded proteins to prolong virion production^15–17^. Members of the BCL-2 family of genes play pro-death and pro-survival roles in modulating apoptosis^18^, which may affect not only viral production but also tissue damage and immune responses. We therefore investigated how the expression of BCL-2 family genes changes in response to SARS-CoV-2 infection within a recently reported study^36^. We first confirmed that productive SARS-CoV-2 infection of lung cell lines Calu-3 and A549 (both WT and ACE2 overexpressing) potently activates the unfolded protein response as evidenced by the upregulation of canonical UPR-associated genes *ATF4* and *DDIT3* (C/EBP homologous protein [CHOP]) (Figure 5A, S3A). Furthermore, active infection also induced expression of pro-apoptotic BIM (*BCL2L11*) and especially the endogenous MCL-1 inhibitor protein Noxa (*PMAIP1*). Upregulation of Noxa is sufficient to induce apoptosis in MCL-1 dependent cells^37,38^ and enhances dependence on other pro-survival proteins such as BCL-2 or BCL-X_L_^39,40^.

**Figure 5:**
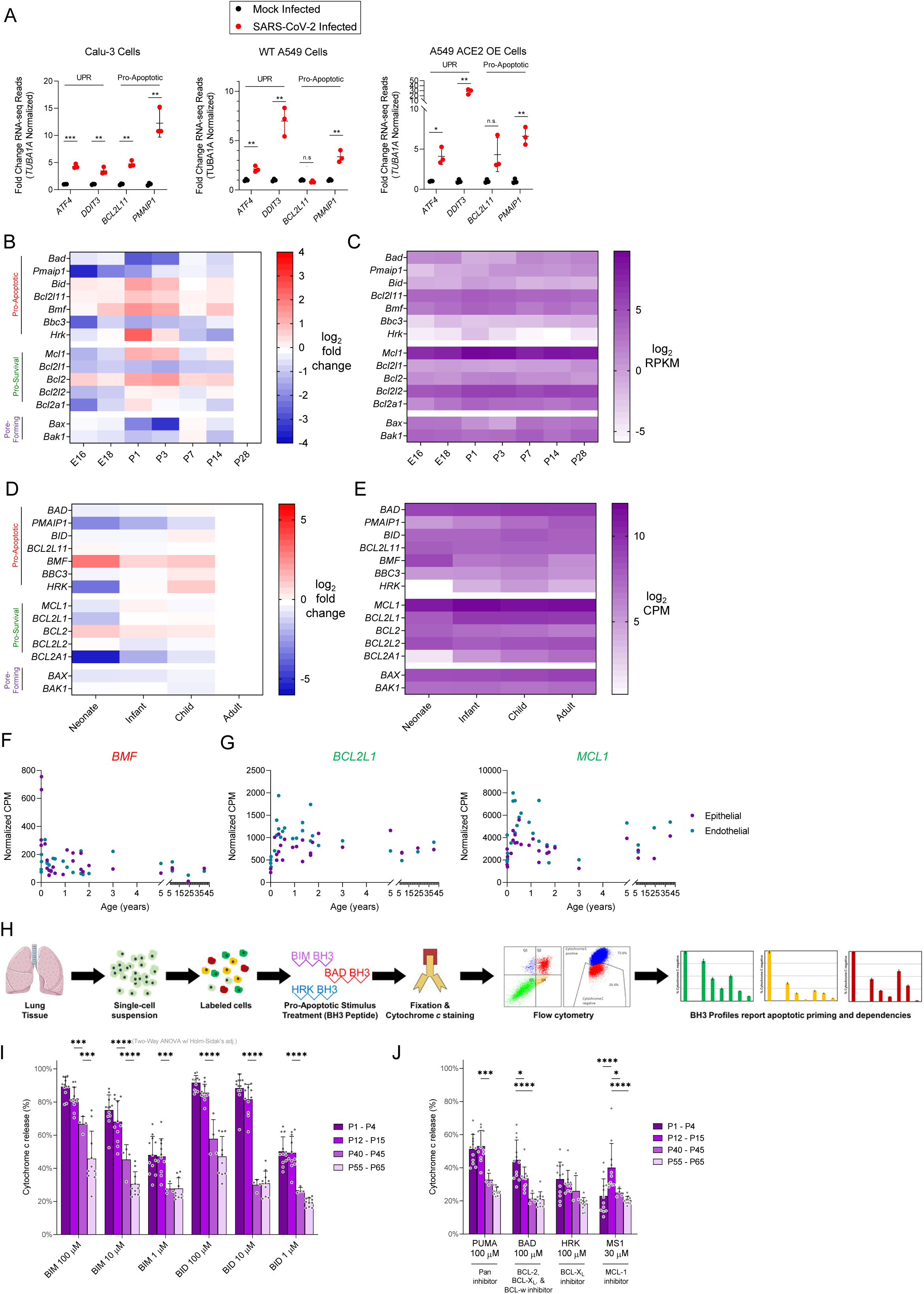
Changing expression of cell death-associated genes may modulate outcomes in children versus adults. (A) RNA sequencing data for selected unfolded protein response and cell death genes in mock and SARS-CoV-2 infected cell lines (B, D) Changes in lung epithelial cell expression of mouse (B) and human (D) Bcl-2 family genes relative to P28 or adult levels. (C, E) Absolute expression levels of Bcl-2 family genes as measured by RNA sequencing of sorted lung epithelial cells in mouse (B) and human (E). (F, G) RNA sequencing data showing pro-apoptotic *BMF* (F) and pro-survival *BCL2L1* and *MCL1* (G) expression in FACS-sorted cell types from human lung. (H) Schematic showing protocol for BH3 profiling assay. (I, J) Apoptotic priming of mouse lung tissue as measured by BH3 profiling. Data represent percentage cytochrome c release upon application of (I) activator or (J) sensitizer BH3 peptides. Comparison between groups was conducted by two-way ANOVA. (*, p<0.05; ***, p<0.001; ****, p<0.0001)

We next assessed how BCL-2 family genes were regulated during postnatal lung maturation to potentially modulate cell fate in response to SARS-CoV-2 infection. In LGEA RNA-seq data, pro-apoptotic genes *Bcl2l11* (encoding BIM) and *Bmf* were both highly expressed in young mouse lung tissue; both peaked at P1 and declined over the remaining timepoints until P28 (late juvenile, early adult) (Figure 5B-C, S2B). Among pro-survival BCL-2 family members, *Mcl1* was most highly expressed, and similarly peaked at P1, as did the more moderately expressed *Bcl2* (Figure S2C). In contrast, expression of *Bcl2l1* (coding for pro-survival BCL-X_L_ protein) was low in P1 mice but increased by P28. *Bax* and *Bak1*, which encode pore-forming proteins that trigger the apoptotic cascade when pro-apoptotic Bcl-2 family members overwhelm pro-survival family members, were expressed at levels likely sufficient to allow apoptosis execution at all timepoints (Figure S2D).

We next tested whether similar expression changes in BCL-2 family genes would be evident in human lung tissue. We examined BCL-2 family RNA-seq data in epithelial cells within the LungMAP database and found that expression patterns were largely consistent with mouse data. Pro-apoptotic *BCL2L11* and *BMF* were again expressed at increased levels in early life (neonates) and reduced in adult lung tissue while pro-survival *MCL1, BCL2A1* (encoding BFL-1/A1) and *BCL2L1* (BCL-X_L_) increased with age (Figure 5D-G, S2E-G). These changes, consistent with a decrease in apoptotic priming over time, were observed predominantly in epithelial and endothelial cells of the human lung.

These gene expression patterns suggested that young lung tissue would be more prone to undergoing apoptotic cell death. We tested this at the functional level using BH3 profiling, an assay that tests mitochondrial sensitivity to titrated doses of apoptosis-inducing peptides (Figure 5H, S2H)^41^. We found that lung epithelial cells (EPCAM^+^) are highly primed for apoptosis at early age as evidenced by higher levels of cytochrome c release in response to pro-apoptotic BIM or BID BH3 peptides in young (P1-P15) versus older lung tissue (Figure 5I). Importantly, adult lung epithelial cells were more resistant to apoptosis, but the pathway remained intact as indicated by continued sensitivity to higher concentrations of BH3 peptides. Interestingly, BH3 profiling also demonstrated that lung epithelial cells in the youngest mice were dependent on BCL-2 and/or BCL-X_L_ for survival but that this dependence switched to MCL-1 in the P12-P15 animals (Figure 5J). Combined with results from cells actively infected with SARS-CoV-2 (Figure 5A-B), these data suggest that infection of lung epithelial cells would trigger apoptosis more quickly and readily in the young lung than in the adult to limit further virion production. Indeed, apoptotic resistance of infected cells has been previously shown to increase virion production, infection severity and host mortality^12,42–45^.

While impairment of lung function is a major source of morbidity and mortality for COVID-19 patients, severe damage to the cardiovascular, renal and gastrointestinal systems by SARS-CoV-2 is also evident in patients with poor outcomes^46^. We therefore compared the expression of SARS-CoV-2 cell entry genes across major organs and found that *ACE2* expression in human lung was low, and *TMPRSS2* expression was high, relative to other organs (Figure 6A) – this is consistent with our immunofluorescence analysis showing cell type-specific lung expression of ACE2 at most ages. Throughout the respiratory system, *TMPRSS2* and *ACE2* were broadly expressed and, consistent with previous reports, *ACE2* expression was particularly high in the nasopharynx^11,47^. Also consistent with previous reports^48^, *ACE2* expression was high in several extrapulmonary tissues, including human testis, kidney, and GI tract – several of these were confirmed at the protein level by mass spectrometry^49^ (Figure 6B).

**Figure 6:**
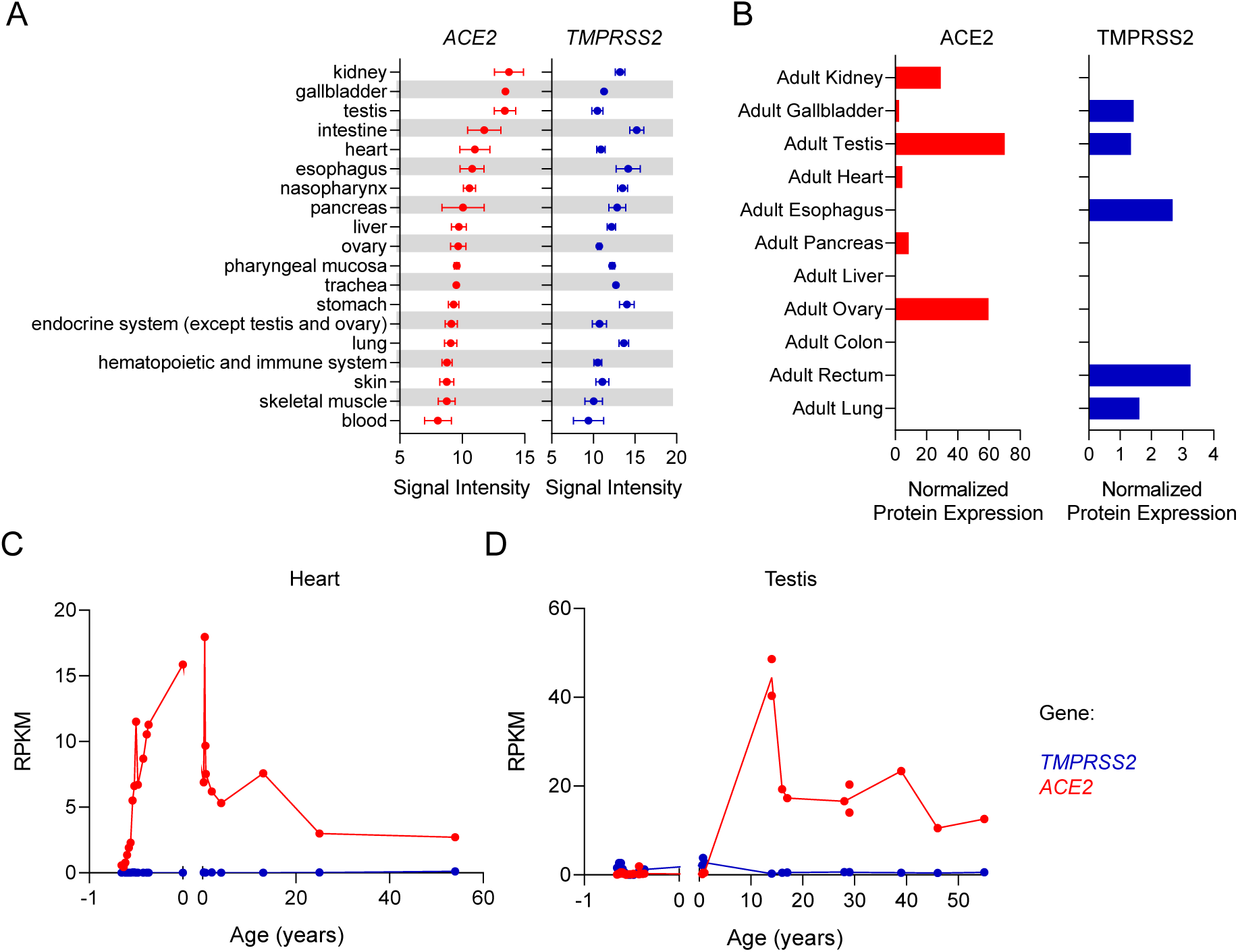
*ACE2* and *TMPRSS2* are dynamically regulated in extrapulmonary human tissues. (A) Microarray expression of *ACE2* and *TMPRSS2* in selected categories of tissue as measured using the Affymetrix Human Genome U133 Plus 2.0 array. (B) Human expression of ACE2 and TMPRSS2 in selected tissues as measured by mass spectrometry (C, D) RNA sequencing data showing age-dependent expression of *ACE2* and *TMPRSS2* in human heart (C) and testis (D) tissue.

Finally, we extended our age-dependent gene expression analysis to additional human tissues to examine how *ACE2* and *TMPRSS2* levels might change over lifespan. While cardiac tissue expressed *ACE2* throughout prenatal and postnatal life, expression levels were particularly high in early childhood and then declined with age (Figure 6C)^50^. It is unclear whether this may be linked to the increasing evidence of cardiac dysfunction in young children diagnosed with COVID-19 (whether via the enzymatic function of ACE2 or its role as a viral entry receptor)^51,52^. A similar pattern was observed in testis tissue (Figure 6D). Little or no *ACE2* expression was detected across lifespan in the brain or liver (Figure S4A-C); limited postnatal expression data was available for kidney and ovary tissue, but available timepoints showed moderate expression in the former and low expression in the latter (Figure S4A-E). Proteomic and genomic measurements were mismatched for some tissues (e.g. ovary), suggesting potential posttranscriptional regulation of ACE2 levels.

## DISCUSSION

Our studies provide evidence that ACE2, TMPRSS2 and apoptotic programs are dynamically regulated by age and cell type in the lung and correlate with severity of COVID-19 disease. Based on our study and our current understanding of SARS-CoV-2 infection, it is plausible that increased expression of ACE2 in airway and alveolar epithelial cells in infancy and old age contributes to the relative severity of COVID-19 symptoms in these populations (Tables 2 and 3). Furthermore, the strong upregulation of ACE2 in late adulthood across multiple cell types in the lung increases the number of cells that can potentially be infected by SARS-CoV-2. ACE2 expression on fragile AT1 cells could be especially problematic given their roles in barrier function and gas exchange. In severe cases of COVID-19, leakage of blood plasma and interstitial fluid into the airspace can occur, along with severe inflammation and hyperproliferation of AT2 cells^46,53^; rapid loss of infected AT1 cells would be expected to promote all these responses.

**Table 3:** Age-dependent regulation of ACE2, TMPRSS2 and apoptosis in human lung

| Human age: |  | 0-30 days<br>(Newborn) | 1-12<br>months<br>(Infant) | 1-17 years<br>(Child) | 18-45<br>years<br>(Adult) | 45-64<br>years<br>(Late<br>Adult) | 65-74<br>years<br>(Elderly) | 75+<br>(Advanced<br>Age) |
| --- | --- | --- | --- | --- | --- | --- | --- | --- |
| ACE2 mRNA<br>Lung Epithelial |  | Low | Moderate/<br>High | Moderate | Moderate | Not<br>available | Not<br>available | Not<br>available |
| ACE2 mRNA<br>Lung Endothelial |  | None | None | None | Low | Not<br>available | Not<br>available | Not<br>available |
| ACE2 in<br>Lung Cells | All (DAPI+) | Not<br>available | High | Low,<br>variable | Moderate,<br>variable | Moderate,<br>variable | High,<br>variable | High,<br>variable |
|  | AQP3+ | Not<br>available | Not<br>available | Moderate,<br>variable | Moderate,<br>variable | Moderate,<br>variable | Moderate,<br>variable | Moderate,<br>variable |
|  | PDPN+ | Not<br>available | Not<br>available | Low,<br>variable | Moderate,<br>variable | Moderate,<br>variable | High,<br>variable | High,<br>variable |
| TMPRSS2 mRNA<br>Lung Epithelial |  | None | Moderate | Low | Low | Not<br>available | Not<br>available | Not<br>available |
| TMPRSS2 mRNA<br>Lung Endothelial |  | None | None | None | Low | Not<br>available | Not<br>available | Not<br>available |
| Apoptotic Priming |  | Not<br>available | Not<br>available | Not<br>available | Not<br>available | Not<br>available | Not<br>available | Not<br>available |
| BCL-2 Family Gene<br>Expression | Pore-Forming | Moderate | Moderate | Moderate | Moderate | Not<br>available | Not<br>available | Not<br>available |
|  | Pro-Apoptotic | Moderate | Moderate | Moderate | Low | Not<br>available | Not<br>available | Not<br>available |
|  | Pro-Survival | Moderate | Moderate | Moderate | High | Not<br>available | Not<br>available | Not<br>available |

Our study also demonstrates that the apoptotic priming of young lung tissue is higher relative to adults; this has been previously associated with decreased virion production due to earlier induction of apoptosis^15,16^. Our results suggest that this mechanism, along with reduced ACE2 expression, may reduce COVID19 disease severity in children.

It is important to note that our study has limited racial diversity among donors of the TMA lung tissue. Only Asian individuals were represented in these TMAs, preventing analysis of racial or ethnic differences in ACE2 expression. However, given the substantial impact of social determinants of health on variable COVID19 mortality rates among racial groups in the U.S.^54^, it is perhaps unlikely that ACE2 expression is a major contributor to these racial disparities.

Our results have important implications for the development of therapeutic approaches to COVID19. One therapeutic under investigation is human recombinant soluble ACE2 (hrsACE2)^55^; our results support the feasibility of this approach, but suggest that with increasing age in patients, the number of available receptors for SARS-CoV-2 continually increases, perhaps necessitating increasing doses of hrsACE2 to “compete away” viral particles from host cell receptors. It remains unclear how the enzymatic activity of hrsACE2 might affect patient physiology and antiviral responses via effects on the renin-angiotensin system. Our findings also suggest a potential therapeutic approach focused on cell death responses to infection, wherein apoptotic priming in adult lung tissue would be modulated to match that in pediatric lung. This approach, which could involve administration of BH3 mimetics (small molecule BCL-2 family inhibitors^56^) systemically or directly to lung tissue via inhalation, would be expected to reduce virus replication in adults as infected, stressed cells would undergo apoptosis earlier. Given the potential role of apoptotic cell death in suppressing inflammation, such a treatment might further reduce negative outcomes in infected patients by immune-mediated mechanisms^57^. Finally, the use of drugs that inhibit the activity of UPR proteins such as PERK or IRE1 may impair ER stress responses, increase apoptotic signaling and accelerate the commitment to apoptosis in infected cells to reduce virion production^16,58^. However, in each of these cases, the potential tissue damaging effects of apoptosis promotion require careful consideration

It is important to note that initial infection and cell death are only two of an extensive set of factors influencing disease course in COVID-19 patients. Immune response^36,59^, host genetics^60^, environmental factors^8^, and therapeutic interventions^61,62^ may all play contributing roles in determining infection outcome. Further, due to the dynamic regulation of blood pressure, fluid and electrolytes by the renin-angiotensin-aldosterone system^63^, it is likely that inter- and intra-individual variation in ACE2 expression will also impact disease course, which is consistent with the extreme heterogeneity in lung ACE2 expression we observed. Despite this, our discovery of the age dependent regulation of ACE2, TMRPRSS2 and apoptosis sensitivity in the lung shed light on potential determinants of disease severity in highly susceptible individuals.

## METHODS

### Animal care and use

Mouse tissue immunofluorescence experiments described in these studies were approved by the Johns Hopkins University Animal Care and Use Committee (Protocol Number: M019M332) and were performed according to the Guide for the Care and Use of Laboratory Animals of the National Institutes of Health. Male and female C57BL/6J mice were obtained from the Jackson Laboratory (#000664, Bar Harbor, ME) and bred and housed in the Johns Hopkins animal facility.

For BH3 profiling, cohorts of mice were housed and bred in a colony in accordance with the policies and regulations set forth by the Harvard TH Chan School of Public Health’s IACUC, under protocol 5245. All animal experiments were approved by IACUC under HSPH protocol #IS00001059-3. Black 6 mice, strain C57BL/6J, (WT) (Jackson Laboratories) were used for tissue collection.

### Human tissue samples

Human tissue microarrays were obtained from US Biomax, Inc. (Derwood, MD). Arrays LCN241 and LC2086a were selected to represent the broadest possible age range of donors and overlapping donors from the two cores were excluded from analysis of LC2086a. Properties of the donors are shown in Supplemental Figure 4. Morphology of the analyzed cores was examined by a pathologist for signs of lung disease; the pathologist’s analysis is shown in Supplemental Figure 4 and cores showing evidence of hemorrhage were excluded to avoid errors in quantification arising from the autofluorescence of red blood cells in the stained cores.

Human infant lung samples were obtained and processed at autopsy from either patients with necrotizing enterocolitis or age-matched infants that died from unrelated conditions that did not affect the lungs, with approval from the University of Pittsburgh Institutional Review Board (CORID No. 491) and in accordance with the University of Pittsburgh anatomical tissue procurement guidelines. All samples were de-identified via an independent honest broker assurance mechanism (Approval #: HB#043) and transferred to Johns Hopkins University under the guidance of MTA approval (JUH MTA # A26558) for analysis.

### Analysis of public gene expression databases

Mouse and human microarray data were analyzed using Genevestigator (NEBION, Zurich, Switzerland). Mouse data were filtered to exclude non-wild type genetic backgrounds and experimental treatments. The remaining lung gene expression data were grouped by age and exported to generate plots. For human microarray data, samples were filtered to exclude disease conditions or drug treatments. The remaining data were grouped by anatomy. Some exported groups included broader (e.g. organ system) or narrower (e.g. organ, organ substructure) classifications; all the sample groups plotted were mutually exclusive.

RNA-seq data was obtained from the LGEA^32^ and LungMAP^33^ databases, with corresponding protein levels in extrapulmonary tissues confirmed using the Human Proteome Map^49^ database. RNA sequencing data from SARS-CoV-2 infected cell lines was obtained from GEO accession GSE147507^36^.

### Single cell RNA sequencing

Previously published^73,74^ single cell RNA sequencing datasets were analyzed for expression of genes of interest. Time course data were from accession GSE119228 and 4 month data were from accession GSE121611. Processing was performed using the Seurat package in R^75^.

### Immunofluorescence sample preparation and imaging

Immunofluorescent staining of lung tissues was performed on 4% paraformaldehyde-fixed 5 μm-thick paraffin sections. The sections were first warmed to 56°C in a vacuum incubator (Isotemp Vacuum Oven, Fisher Scientific) then washed immediately twice in xylene, gradually re-dehydrated in ethanol (100%, 95%, 70%, water), and then processed for antigen retrieval by microwave heating (1000 watt, 6 minutes) in citrate buffer (10mM, pH6.0). Samples were then washed with PBS, blocked with 1% BSA/5% donkey-serum (1 hour, room temperature), then incubated overnight at 4°C with primary antibodies (1:200 dilutions in 0.5% BSA). The following day, samples were washed 3 times with PBS and incubated with appropriate fluorescent-labeled secondary antibodies (1:1000 dilution in 0.5% BSA, Life Technologies Inc) and the nuclear marker DAPI (Biolegend). Slides were mounted using Gelvatol (Sigma-Aldrich) solution prior to imaging. Initial imaging was carried out using a 40x/1.3NA objective lens on a Nikon Eclipse Ti Confocal microscope (Nikon, Melville, NY). Pixel sizes were 0.15 microns and z step size of 5 microns. High-resolution imaging was then conducted on the identical slides with a Fluoview 1000 (Olympus Waltham, MA). Antibodies used for mouse lung immunostaining are listed in Table 4. Human TMA samples were imaged on a widefield Rarecyte Cytefinder slide scanner with a 20x/0.75NA objective lens with pixel size 0.65 microns/pixel. The excitation and emission filters were DAPI (ex: 395/25, em: 438/26), FITC (ex: 485/25, em: 522/20), Cy3 (ex: 555/20, em: 590/20), and Cy5 (ex: 651/11, em: 692/44). Exposure times were adjusted to maximize dynamic range and eliminate saturation. High resolution representative images from selected cores were acquired using a Zeiss LSM 880 confocal microscope.

**Table 4:**
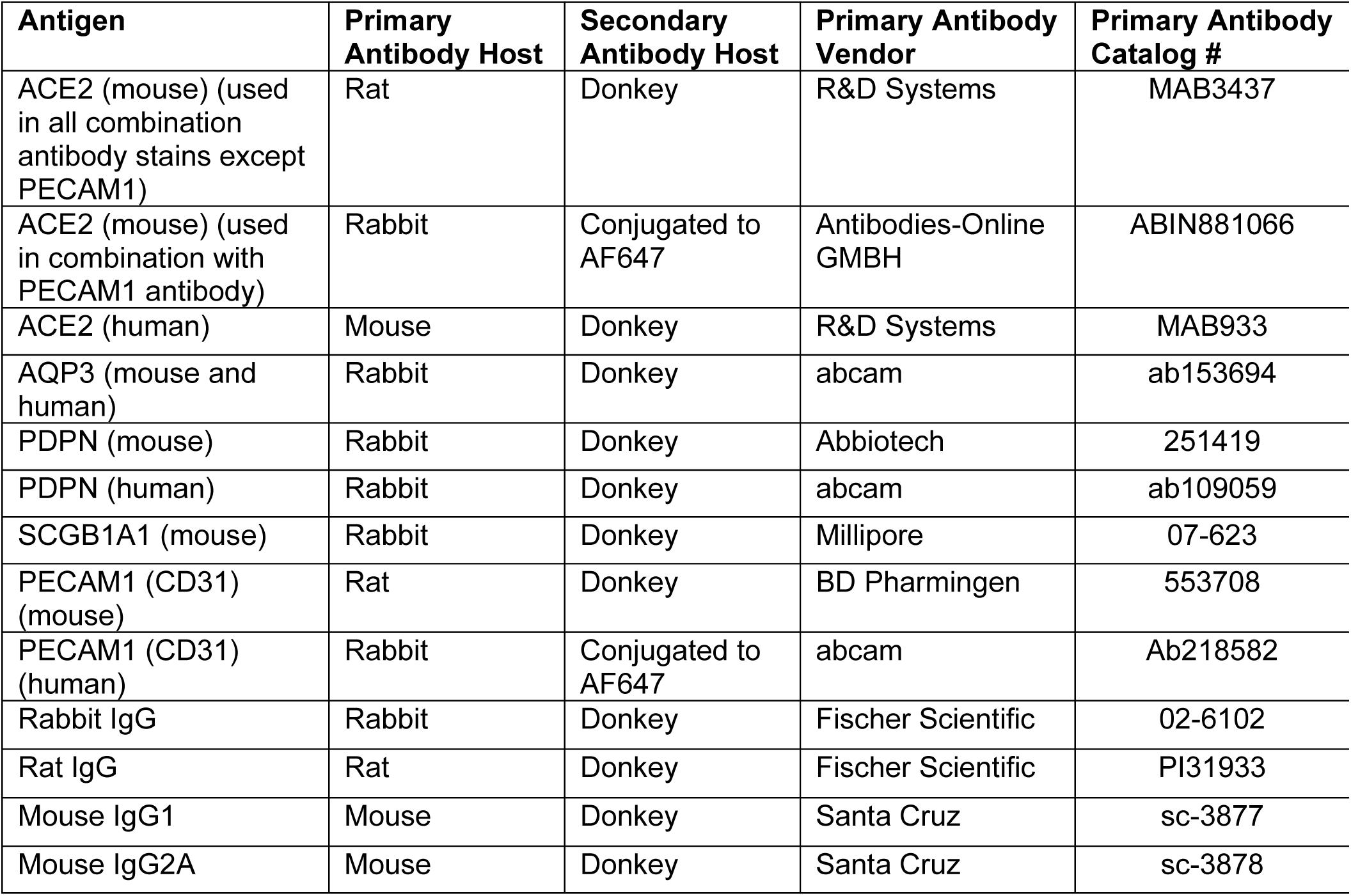
Immunofluorescence antibodies

### Image quantification

For quantification of immunofluorescence images of mouse samples, image datasets were saved as .nd2 files and analyzed as maximum intensity projections since tissue samples had a thickness of one cell layer. Cells were segmented into individual objects based on the DAPI channel. Due to the fact that nuclei boundaries are ambiguous and hard to identify in tissue, we applied a preprocessing semantic segmentation step using a UNet model^76^ trained on hand annotated DAPI-stained mouse and human tissue (https://github.com/HMS-IDAC/UnMicst). The architecture of the implemented model was similar to Saka et al. (2019)^77^. Briefly, the model was trained to recognize and generate probability maps for nuclei contours, nuclei centers, and background pixels. Cellprofiler^78^ was then used to further segment nuclei based on the nuclei center probability maps made by the UNet model.

In Cellprofiler, nuclei were identified using the Identify Primary Objects module where clumped cells were identified based on shape. Next, the corresponding whole cell region was obtained using the Identify Secondary Objects and dilating by 4 pixels since a common cytoplasm channel was absent from the experiment. The cytoplasmic regions were then obtained using Identify Tertiary Objects. Before acquiring measurements, the green, red and Cy5 channels were background subtracted by sampling the background intensities in the lower quartile range. The background subtracted median intensities were then measured on a per-cell basis and exported to a csv file separated by nuclei, cytoplasm, and cell mask regions with entire cell mask region being used for further analysis. Segmentation mask overlays were also saved and inspected to evaluate the quality of segmentation. Image quantification values were reported as background-subtracted median ACE2 intensities in either all cells (DAPI^+^) or in cells positive (above background) for indicated markers.

For quantification of the human TMA samples, images were acquired as separate overlapping tiles from a slidescanner. These tiles were stitched using ASHLAR, a novel stitching and registration algorithm (https://github.com/labsyspharm/ashlar). Other custom scripts were used to 1) systematically separate the stitched image into organized individual tissue cores (Coreograph; https://github.com/HMS-IDAC/UNetCoreograph), preprocess the image to identify cells and suppress artefacts based on the DAPI channel (UnMicst; https://github.com/HMS-IDAC/UnMicst), and segment cells into individual objects (S3segmenter; https://github.com/HMS-IDAC/S3segmenter). Upon closer inspection, we noticed that the staining quality of DAPI on the tissue cores was variable despite staining all cores under the same conditions. It is known that DNA deteriorates with age of sample causing lower binding of DAPI. This causes the signal-to-background ratio to be low where the nuclei are obfuscated by tissue autofluorescence and finally leading to poor nuclei segmentation. We therefore inspected each core and only retained cores that had prominent DAPI staining above background. From these remaining cores, background subtracted mean and median intensity measurements were obtained on a single-cell basis and exported to a csv file. Because running a large number of cores manually would be tedious, prone to errors, and require significant computational resources, all processes (stitching, core separation, preprocessing, single cell segmentation, and quantification) were incorporated into a unified automated pipeline (mcmicro; https://github.com/labsyspharm/mcmicro) that processed all samples in parallel on a high performance cluster. The background subtracted median intensities were then measured on a per-cell basis and exported to a csv file separated by nuclei, cytoplasm, and cell mask regions with entire cell mask being used for further analysis. Cores containing fewer than 100 cells of a given type were excluded from analysis. Segmentation mask overlays were also saved and inspected to evaluate the quality of segmentation. Image quantification values were reported as background-subtracted ACE2 median intensities in either all cells (DAPI^+^) or in cells positive (above background) for indicated markers.

### BH3 profiling

Lung samples from mice of different ages (P0-P62) were dissociated into a single cell suspension using the Papain Dissociation System (Worthington Biochemical Corporation) with a modified protocol. Briefly, 50-100g lung samples were roughly chopped and submerged in 500μL of EBSS with 20 units/ml papain and 0.005% DNase and incubated at 37°C with frequent agitation for 15 minutes. Samples were placed on ice and triturated with cut 1ml pipette, and were left to settle for 2-5 minutes before the cloudy cell suspension was transferred to new tubes and centrifuged at 200g for 5 minutes at 4°C. The resulting pellet was resuspended in EBSS with 0.005% DNase, 1mg/ml bovine serum albumin and 1mg/ml Ovomucoid protease inhibitor. This suspension was layered on top of a solution of EBSS with 10mg/ml bovine serum albumin and 10mg/ml Ovomucoid protease inhibitor to create a discontinuous density gradient, and then centrifuged at 72g for 6 minutes at room temperature. Supernatant was discarded and the pellet was resuspended in 100µl FACS Stain Buffer (2% FBS in PBS) with 1µl anti-CD45-APC/Cy7 (clone 30-F11, BioLegend) and 1µl anti-EPCAM-AlexaFluor488 (clone G8.8, BioLegend).

Cells were stained on ice for 25 minutes away from light, then centrifuged at 200g for 5 minutes and subjected to flow cytometry-based BH3 profiling as previously described^41^. Briefly, cells were treated with activator or sensitizer BH3 peptides (New England Peptide) for 60 minutes at 28°C in MEB (10 mM HEPES pH 7.5, 150 mM mannitol, 50 mM KCl, 0.02 mM EGTA, 0.02 mM EDTA, 0.1% BSA, 5 mM succinate) with 0.001% digitonin. Peptide sequences are as follows: BIM (Ac-MRPEIWIAQELRRIGDEFNA-NH2), BID (Ac-EDIIRNIARHLAQVGDSMDRY-NH2), PUMA (Ac-EQWAREIGAQLRRMADDLNA-NH2), BAD (Ac-LWAAQRYGRELRRMSDEFEGSFKGL-NH2), HRK (Ac-WSSAAQLTAARLKALGDELHQ-NH2), and MS1 (Ac-RPEIWMTQGLRRLGDEINAYYAR-NH2). After peptide exposure, cells were fixed in 2% paraformaldehyde for 15 minutes which was then neutralized by addition of N2 buffer (1.7 M Tris base, 1.25 M glycine, pH 9.1). Cells were stained overnight with DAPI (1:1000, Abcam) and anti-Cytochrome c-AlexaFluor647 (1:2000, clone 6H2.B4, Biolegend) in a saponin-based buffer (final concentration 0.1% saponin, 1% BSA) and then analyzed by flow cytometry. Cytochrome c release in response to BIM treatment was measured on an Attune NxT flow cytometer (Thermo Fisher Scientific). Data for EPCAM+ lung epithelial cells collected from the DAPI+/CD45-population.

## ACKNOWLEDGEMENTS

We thank the members of our labs for their comments and suggestions on this work, especially Kaitlyn Webster for assistance with image quantification. We thank Olesja Popow (Dana Farber Cancer Institute) for providing training data for the UNet model and Zoltan Maliga (Harvard Program in Therapeutic Science) for imaging the human TMA samples and advising on staining protocols. Imaging experiments were supported in part by the Emory University Integrated Cellular Imaging Microscopy Core. This work was supported by funding from the HSPH Dean’s Fund for Scientific Advancement (K.A.S.), the HSPH National Institute for Environmental Health Sciences (NIEHS) Center (K.A.S., J.S.), R00CA188679 (K.A.S.), R01CA248565 (K.A.S.), R21AI149321 (H.J.), and R01AI148446 (H.J.), Harvard Center for Cancer Systems Pharmacology, U54CA225088 (C.Y.), Precancer Atlas Program, U2CCA233262 (C.Y.), National Cancer Institute, U2CCA233280 (C.Y.), and DARPA Biostasis, W911NF-19-2-0017 (C.Y.).

## AUTHOR CONTRIBUTIONS

K.S. and Z.I. conceived the study. Z.I., C.Y., B.A.C., B.D., E.G., D.B., and K.S. analyzed data. Z.I., B.D., H.J., and G.N.J conducted imaging experiments. Z.I., J.S., and C.F. conducted BH3 profiling experiments. C.S., D.J.H., and H.J. provided samples for analysis. L.K. conducted pathological analysis of samples. Z.I. and K.S. wrote the manuscript. All authors edited and approved the final manuscript.

## COMPETING INTEREST DECLARATION

The authors declare no competing interests

**Supplementary Figure S1:**
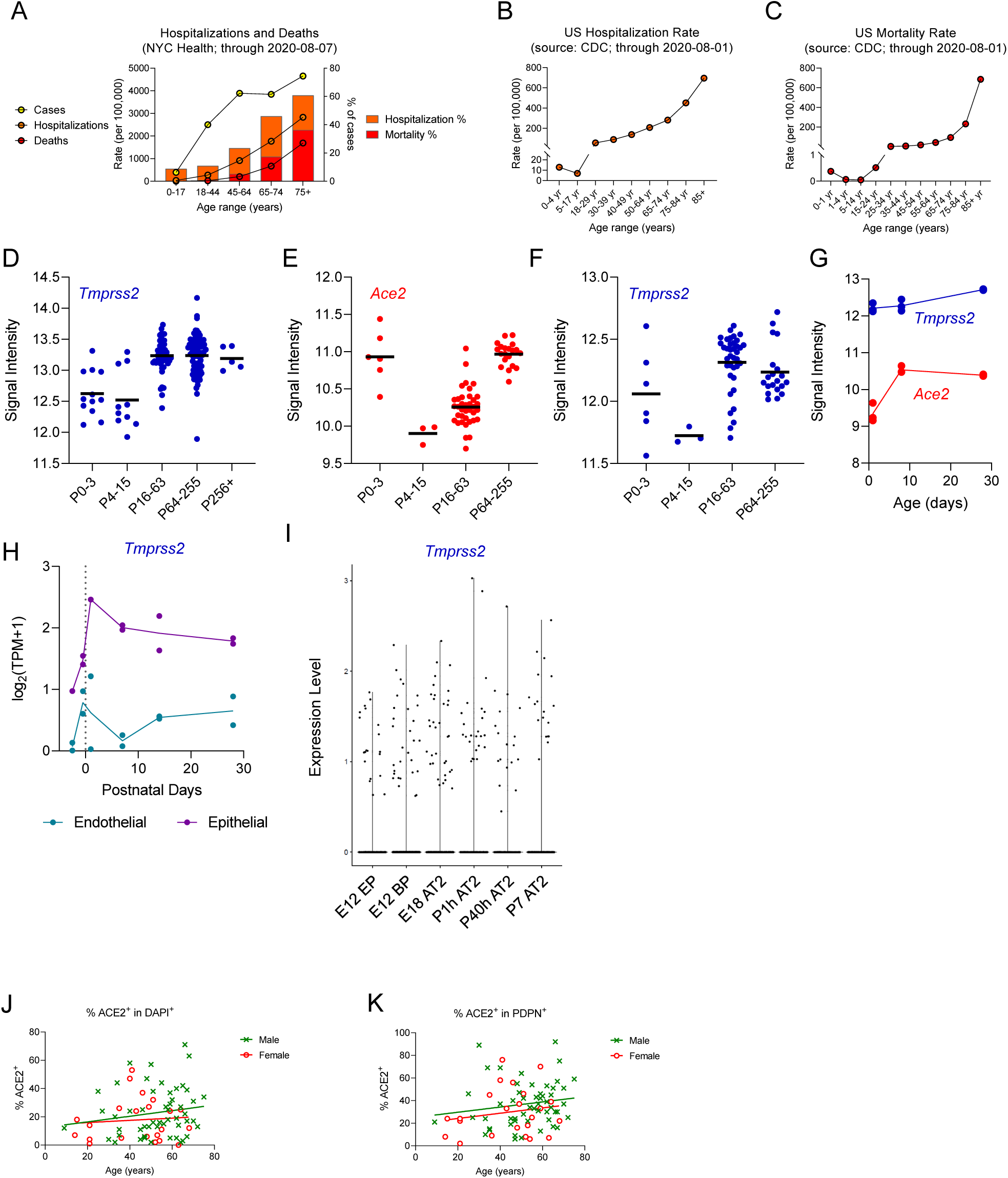
(A) COVID-19 case statistics for New York City as of 08/08/2020. Case (SARS-CoV-2+ tests), hospitalization and death rates are presented per 100,000 population (left axis) or as percentage of cases (right axis). (B-C) COVID-19 hospitalization (B) and mortality (C) rate per 100,000 population in the United States as of 08/01/2020. (D-F) Collected mouse microarray lung gene expression data. (D) Collected *Tmprss2* expression data from experiments conducted using the Affymetrix Mouse Genome 430 2.0 array. (E, F) Collected *Ace2* (E) and *Tmprss2* (F) expression data from experiments conducted using the Affymetrix Mouse Gene ST 1.0 array. (G) Microarray expression data for *Ace2* and *Tmprss2* in whole mouse lung. (H) RNA sequencing data showing *Tmprss2* expression in FACS-sorted cell types from mouse lung. (I) Single cell RNA sequencing showing *Tmprss2* expression in AT2 cells and their precursors in mouse lung at the indicated pre- and postnatal timepoints. (J-K) Scatter plots comparing percentage of all nucleated (J) or alveolar type 1 cells (K) that are positive for ACE2 in male versus female normal lung specimens.

**Supplementary Figure S2:**
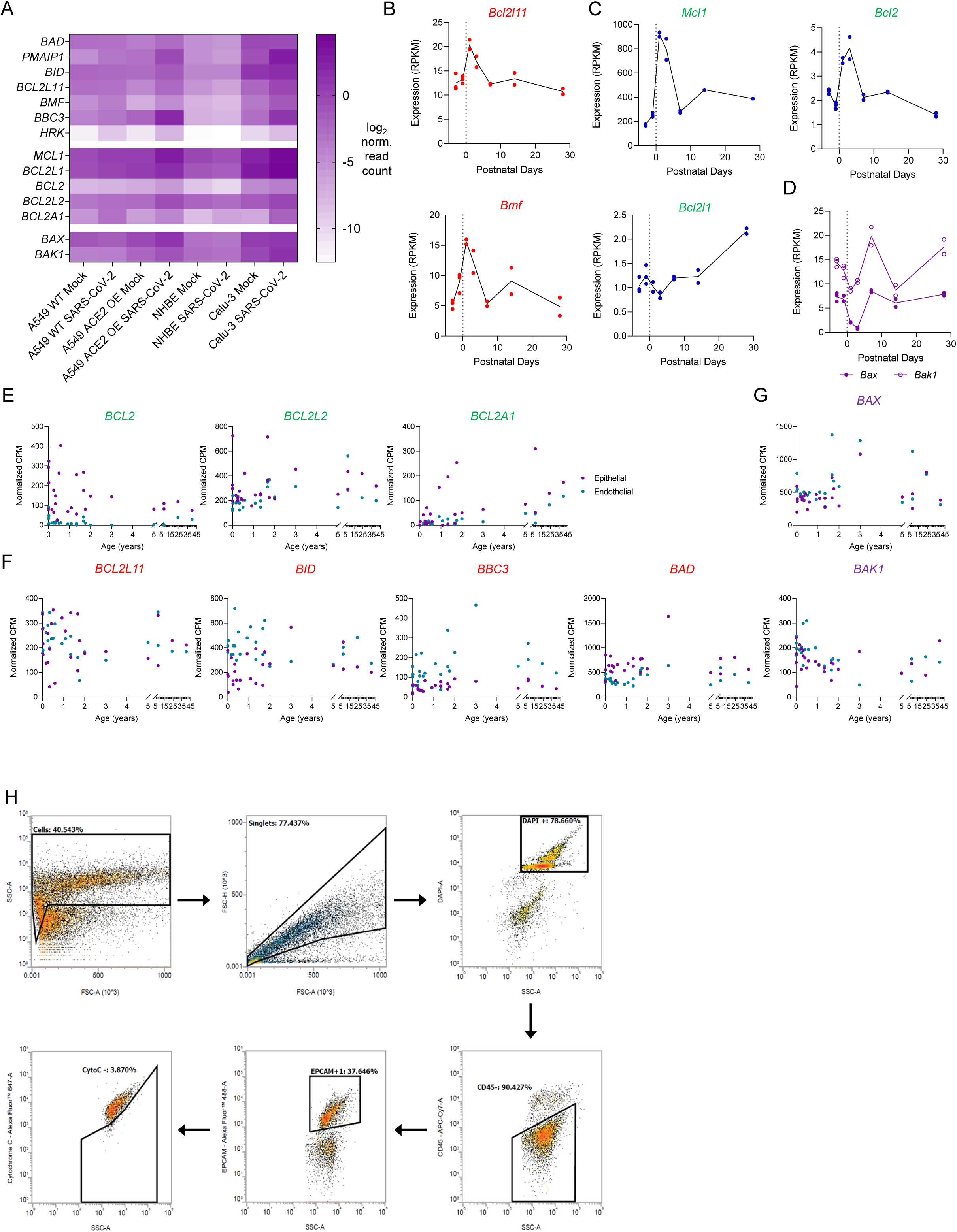
(A) RNA sequencing data for cell death genes in mock and SARS-CoV-2 infected cell lines (B-D) RNA sequencing data showing whole lung expression of the indicated genes: pro-apoptotic *Bcl2l11* and *Bmf* (B) pro-survival *Mcl1, Bcl2l1*, and *Bcl2* (C), and pore-forming *Bax* and *Bak1* (D). (E-G) RNA sequencing data showing pro-survival (E), pro-apoptotic (F), and pore-forming (G) gene expression in FACS-sorted cell types from human lung. (H) Single-cell BH3 profiling analysis was performed on live single cells, chosen according to their FSC and SSC parameters. Lung epithelial cells were selected as DAPI^+^/CD45^−^/EPCAM^+^ and analyzed for % cytochrome c negative in response to different concentrations of BIM peptide.

**Supplementary Figure S3:**
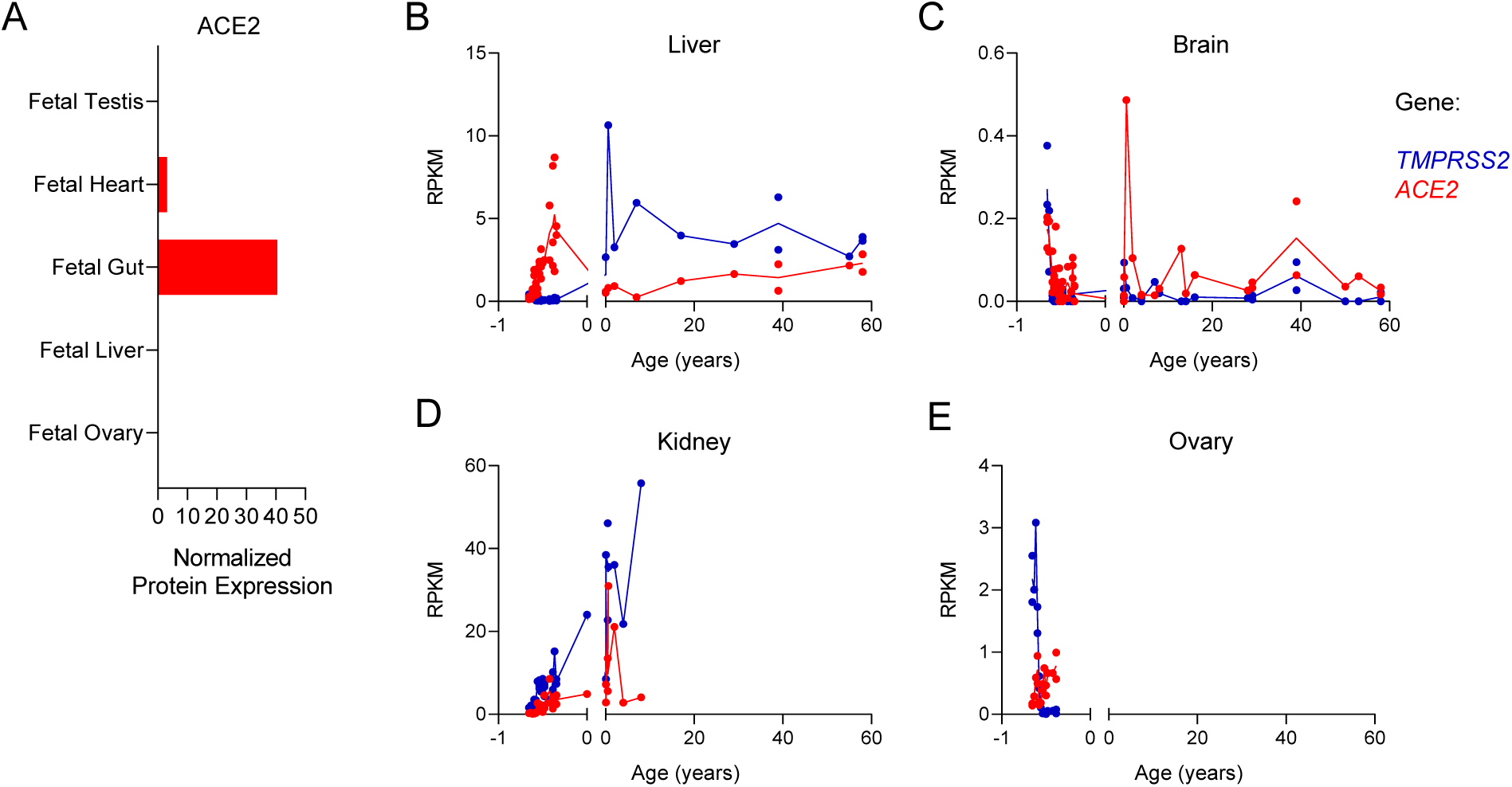
(A) Human expression of ACE2 in selected tissues as measured by proteomic profiling. (B-E) RNA sequencing data showing age-dependent expression of *ACE2* and *TMPRSS2* in human liver (B), brain (C), kidney (D), and ovary (E).

**Supplementary Figure S4:**
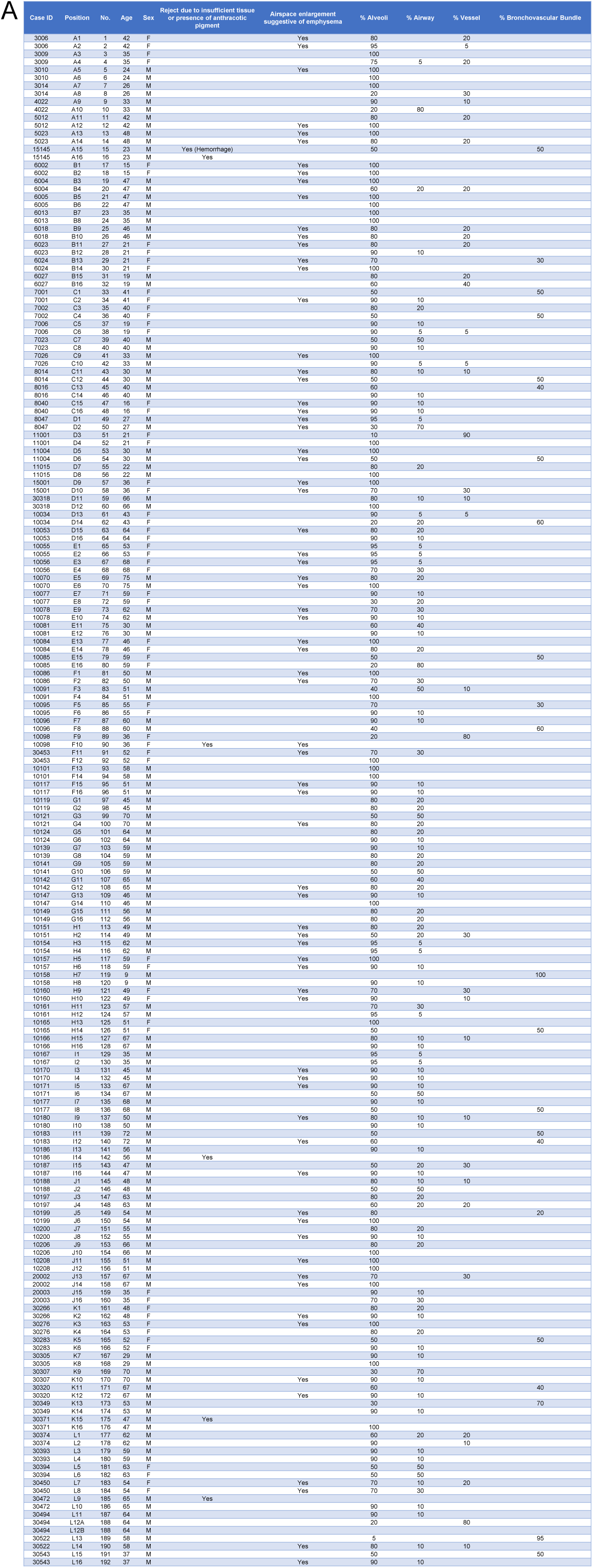
Human lung tissue microarray characteristics. (A) Human lung specimens were stained and examined for tissue composition and indications of unsuitability for analysis.

## REFERENCES

1. Hoffmann, M. et al. SARS-CoV-2 Cell Entry Depends on ACE2 and TMPRSS2 and Is Blocked by a Clinically Proven Protease Inhibitor. Cell 1–10 (2020). doi: 10.1016/j.cell.2020.02.052

2. WHO. Situation reports. (2020).

3. Bialek, S. et al. Coronavirus Disease 2019 in Children — United States, February 12–April 2, 2020. MMWR. Morb. Mortal. Wkly. Rep. 69, 422–426 (2020).

4. Dong, Y. et al. Epidemiological Characteristics of 2143 Pediatric Patients With 2019 Coronavirus Disease in China. Pediatrics (2020). doi: 10.1542/peds.2020-0702

5. Zhu, N. et al. A novel coronavirus from patients with pneumonia in China, 2019. N. Engl. J. Med. 382, 727–733 (2020).

6. Treutlein, B. et al. Reconstructing lineage hierarchies of the distal lung epithelium using single-cell RNA-seq. Nature 509, 371–375 (2014).

7. Jia, H. Pulmonary Angiotensin-Converting Enzyme 2 (ACE2) and Inflammatory Lung Disease. SHOCK 46, 239–248 (2016).

8. Smith, J. C. & Sheltzer, J. M. Cigarette smoke triggers the expansion of a subpopulation of respiratory epithelial cells that express the SARS-CoV-2 receptor ACE2. bioRxiv 2020.03.28.013672 (2020). doi: 10.1101/2020.03.28.013672

9. Zhao, Y. et al. Single-cell RNA expression profiling of ACE2, the putative receptor of Wuhan 2019-nCov. bioRxiv 2020.01.26.919985 (2020). doi: 10.1101/2020.01.26.919985

10. Travaglini, K. J. et al. A molecular cell atlas of the human lung from single cell RNA sequencing. bioRxiv 7191, 742320 (2019).

11. Ziegler, C. G. K. et al. SARS-CoV-2 receptor ACE2 is an interferon-stimulated gene in human airway epithelial cells and is enriched in specific cell subsets across tissues. CellSneakPeak (2020).

12. Danthi, P. Viruses and the Diversity of Cell Death. Annu. Rev. Virol. 3, 533–553 (2016).

13. Orzalli, M. H. & Kagan, J. C. Apoptosis and Necroptosis as Host Defense Strategies to Prevent Viral Infection. Trends in Cell Biology 27, 800–809 (2017).

14. Fung, T. S. & Liu, D. X. Coronavirus infection, ER stress, apoptosis and innate immunity. Front. Microbiol. 5, (2014).

15. Deng, X. et al. Coronavirus nonstructural protein 15 mediates evasion of dsRNA sensors and limits apoptosis in macrophages. Proc. Natl. Acad. Sci. U. S. A. 114, E4251–E4260 (2017).

16. DeDiego, M. L. et al. Severe Acute Respiratory Syndrome Coronavirus Envelope Protein Regulates Cell Stress Response and Apoptosis. PLoS Pathog. 7, e1002315 (2011).

17. Tan, Y.-J., Lim, S. G. & Hong, W. Regulation of cell death during infection by the severe acute respiratory syndrome coronavirus and other coronaviruses. Cell. Microbiol. 9, 2552–2561 (2007).

18. Singh, R., Letai, A. & Sarosiek, K. Regulation of apoptosis in health and disease: the balancing act of BCL-2 family proteins. Nat. Rev. Mol. Cell Biol. 1 (2019). doi: 10.1038/s41580-018-0089-8

19. Sarosiek, K. A. et al. Developmental Regulation of Mitochondrial Apoptosis by c-Myc Governs Age- and Tissue-Specific Sensitivity to Cancer Therapeutics. Cancer Cell 31, 142–156 (2017).

20. Hoffmann, M. et al. SARS-CoV-2 Cell Entry Depends on ACE2 and TMPRSS2 and Is Blocked by a Clinically Proven Protease Inhibitor. Cell 181, 271–280.e8 (2020).

21. Jansing, N. L. et al. Unbiased quantitation of alveolar type II to alveolar type i cell transdifferentiation during repair after lung injury in mice. Am. J. Respir. Cell Mol. Biol. 57, 519–526 (2017).

22. Engel, K. B. & Moore, H. M. Effects of preanalytical variables on the detection of proteins by immunohistochemistry in formalin-fixed, paraffin-embedded tissue. Arch. Pathol. Lab. Med. 135, 537–543 (2011).

23. Olajuyin, A. M., Zhang, X. & Ji, H. L. Alveolar type 2 progenitor cells for lung injury repair. Cell Death Discov. 5, (2019).

24. Pan, H., Deutsch, G. H. & Wert, S. E. Comprehensive anatomic ontologies for lung development: A comparison of alveolar formation and maturation within mouse and human lung. J. Biomed. Semantics 10, 1–21 (2019).

25. Snoeck, H. W. Modeling human lung development and disease using pluripotent stem cells. Dev. 142, 13–16 (2015).

26. Schittny, J. C. Development of the lung. Cell Tissue Res. 367, 427–444 (2017).

27. Gretebeck, L. M. & Subbarao, K. Animal models for SARS and MERS coronaviruses. Current Opinion in Virology 13, 123–129 (2015).

28. Hruz, T. et al. Genevestigator V3: A Reference Expression Database for the Meta-Analysis of Transcriptomes. Adv. Bioinformatics (2008). doi: 10.1155/2008/420747

29. Saini, Y. et al. Gene expression in whole lung and pulmonary macrophages reflects the dynamic pathology associated with airway surface dehydration. BMC Genomics 15, 726 (2014).

30. Cho, H. Y. et al. Targeted deletion of Nrf2 impairs lung development and oxidant injury in neonatal mice. Antioxidants Redox Signal. 17, 1066–1082 (2012).

31. Srisuma, S. et al. Fibroblast growth factor receptors control epithelial-mesenchymal interactions necessary for alveolar elastogenesis. Am. J. Respir. Crit. Care Med. 181, 838–850 (2010).

32. Du, Y. et al. Lung Gene Expression Analysis (LGEA): An integrative web portal for comprehensive gene expression data analysis in lung development. Thorax 72, 481–484 (2017).

33. Ardini-Poleske, M. E. et al. LungMAP: The molecular atlas of lung development program. American Journal of Physiology – Lung Cellular and Molecular Physiology 313, L733–L740 (2017).

34. Gaudette, B. T., Iwakoshi, N. N. & Boise, L. H. Bcl-xL protein protects from C/EBP homologous protein (CHOP)-dependent apoptosis during plasma cell differentiation. J. Biol. Chem. 289, 23629–23640 (2014).

35. Sarosiek, K. A. et al. Efficacy of bortezomib in a direct xenograft model of primary effusion lymphoma. Proc. Natl. Acad. Sci. U. S. A. 107, 13069–74 (2010).

36. Blanco-Melo, D. et al. Imbalanced host response to SARS-CoV-2 drives development of COVID-19. (2020). doi: 10.1016/j.cell.2020.04.026

37. Wensveen, F. M. et al. Apoptosis threshold set by noxa and Mcl-1 after T cell activation regulates competitive selection of high-affinity clones. Immunity 32, 754–765 (2010).

38. Haschka, M. D. et al. The NOXA–MCL1–BIM axis defines lifespan on extended mitotic arrest. Nat. Commun. 6, 6891 (2015).

39. Chen, L. et al. Differential targeting of prosurvival Bcl-2 proteins by their BH3-only ligands allows complementary apoptotic function. Mol. Cell 17, 393–403 (2005).

40. Fraser, C., Presser, A., Sanchorawala, V., Sarosiek, S. & Sarosiek, K. Clonal plasma cells in AL amyloidosis are dependent on pro-survival BCL-2 family proteins and sensitive to BH3 mimetics. bioRxiv (2019).

41. Fraser, C., Ryan, J. & Sarosiek, K. BH3 Profiling: A Functional Assay to Measure Apoptotic Priming and Dependencies. in Methods in Molecular Biology 1877, 61–76 (2019).

42. Austgen, K., Oakes, S., virology, D. G.-J. of & 2012, undefined. Multiple defects, including premature apoptosis, prevent Kaposi’s sarcoma-associated herpesvirus replication in murine cells. Am Soc Microbiol

43. Hui, K., Lam, B., Ho, D., … S. T.-M. cancer & 2013, undefined. Bortezomib and SAHA synergistically induce ROS-driven caspase-dependent apoptosis of nasopharyngeal carcinoma and block replication of Epstein–Barr virus. AACR

44. DeDiego, M., … J. N.-T.-Pl. & 2011, undefined. Severe acute respiratory syndrome coronavirus envelope protein regulates cell stress response and apoptosis. ncbi.nlm.nih.gov

45. Fung, T., microbiology, D. L.-F. in & 2014, undefined. Coronavirus infection, ER stress, apoptosis and innate immunity. frontiersin.org

46. Fox, S. E., Akmatbekov, A., Harbert, J. L., Li, G. & Brown, J. Q. Pulmonary and Cardiac Pathology in Covid-19 : The First Autopsy Series from New Orleans 1) Department of Pathology, LSU Health Sciences Center, New Orleans 2) Pathology and Laboratory Medicine Service, Southeast Louisiana Veterans Healthcare System 3. medRxiv (2020).

47. Wu, C. & Zheng, M. Single-cell RNA expression profiling shows that ACE2, the putative receptor of Wuhan 2019-nCoV, has significant expression in the nasal, mouth, lung and colon tissues, and tends to be co-expressed with HLA-DRB1 in the four tissues. (Preprints, 2020).

48. Gupta, A. et al. Extrapulmonary manifestations of COVID-19. Nat. Med. 26, (2020).

49. Kim, M.-S. et al. A draft map of the human proteome. Nature 509, 575–81 (2014).

50. Cardoso-Moreira, M. et al. Gene expression across mammalian organ development. Nature 571, 505–509 (2019).

51. Qiu, H. et al. Clinical and epidemiological features of 36 children with coronavirus disease 2019 (COVID-19) in Zhejiang, China: an observational cohort study. Elsevier

52. Cui, Y., Tian, M., Huang, D., … X. W.-T. J. of & 2020, undefined. A 55-day-old female infant infected with 2019 novel coronavirus disease: presenting with pneumonia, liver injury, and heart damage. academic.oup.com

53. Madjid, M., Safavi-Naeini, P., Solomon, S. D. & Vardeny, O. Potential Effects of Coronaviruses on the Cardiovascular System: A Review. JAMA Cardiol. 10, 1–10 (2020).

54. Abrams, E. M. & Szefler, S. J. COVID-19 and the impact of social determinants of health. The Lancet Respiratory Medicine 8, 659–661 (2020).

55. Monteil, V., Kwon, H., Prado, P., Hagelkrüys, A. & Wimmer, R. Inhibition of SARS-CoV-2 infections in engineered human tissues using clinical-grade soluble human ACE2. cell.com

56. Sarosiek, K. A. & Letai, A. Directly targeting the mitochondrial pathway of apoptosis for cancer therapy with BH3 mimetics: recent successes, current challenges and future promise. FEBS J. 283, 3523–3533 (2016).

57. Medina, C. B. et al. Metabolites released from apoptotic cells act as tissue messengers. Nature 1–6 (2020). doi: 10.1038/s41586-020-2121-3

58. Fung, T., Liao, Y., virology, D. L.-J. of & 2014, undefined. The endoplasmic reticulum stress sensor IRE1α protects cells from apoptosis induced by the coronavirus infectious bronchitis virus. Am Soc Microbiol

59. To, K. K.-W. et al. Temporal profiles of viral load in posterior oropharyngeal saliva samples and serum antibody responses during infection by SARS-CoV-2: an observational cohort study. Lancet Infect. Dis. 0, (2020).

60. Williams, F. M. et al. Self-reported symptoms of covid-19 including symptoms most predictive of SARS-CoV-2 infection, are heritable. medRxiv 2020.04.22.20072124 (2020). doi: 10.1101/2020.04.22.20072124

61. Horby, P. et al. Dexamethasone for COVID-19-Preliminary Report Effect of Dexamethasone in Hospitalized Patients with COVID-19 – Preliminary Report. medRxiv 2020.06.22.20137273 (2020). doi: 10.1101/2020.06.22.20137273

62. Hansen, J. et al. Studies in humanized mice and convalescent humans yield a SARS-CoV-2 antibody cocktail. Science (80-.). eabd0827 (2020). doi: 10.1126/science.abd0827

63. Burrell, L., Johnston, C., … C. T.-T. in E. & & 2004, undefined. ACE2, a new regulator of the renin–angiotensin system. Elsevier

64. Wentworth, D. E., Gillim-Ross, L., Espina, N. & Bernard, K. A. Mice susceptible to SARS coronavirus. Emerg. Infect. Dis. 10, 1293–1296 (2004).

65. Subbarao, K. et al. Prior Infection and Passive Transfer of Neutralizing Antibody Prevent Replication of Severe Acute Respiratory Syndrome Coronavirus in the Respiratory Tract of Mice. J. Virol. 78, 3572–3577 (2004).

66. Glass, W. G., Subbarao, K., Murphy, B. & Murphy, P. M. Mechanisms of Host Defense following Severe Acute Respiratory Syndrome-Coronavirus (SARS-CoV) Pulmonary Infection of Mice. J. Immunol. 173, 4030–4039 (2004).

67. Roberts, A. et al. A mouse-adapted SARS-coronavirus causes disease and mortality in BALB/c mice. PLoS Pathog. 3, 0023–0037 (2007).

68. Baas, T. et al. Genomic Analysis Reveals Age-Dependent Innate Immune Responses to Severe Acute Respiratory Syndrome Coronavirus. J. Virol. 82, 9465–9476 (2008).

69. Tseng, C.-T. K. et al. Severe Acute Respiratory Syndrome Coronavirus Infection of Mice Transgenic for the Human Angiotensin-Converting Enzyme 2 Virus Receptor. J. Virol. 81, 1162–1173 (2007).

70. Roberts, A. et al. Aged BALB/c Mice as a Model for Increased Severity of Severe Acute Respiratory Syndrome in Elderly Humans. J. Virol. 79, 5833–5838 (2005).

71. Hogan, R. J. et al. Resolution of Primary Severe Acute Respiratory Syndrome-Associated Coronavirus Infection Requires Stat1. J. Virol. 78, 11416–11421 (2004).

72. McCray, P. B. et al. Lethal Infection of K18-hACE2 Mice Infected with Severe Acute Respiratory Syndrome Coronavirus. J. Virol. 81, 813–821 (2007).

73. Cohen, M. et al. Lung Single-Cell Signaling Interaction Map Reveals Basophil Role in Macrophage Imprinting. Cell 175, 1031–1044.e18 (2018).

74. Reyfman, P. A. et al. Single-cell transcriptomic analysis of human lung provides insights into the pathobiology of pulmonary fibrosis. Am. J. Respir. Crit. Care Med. 199, 1517–1536 (2019).

75. Stuart, T. et al. Comprehensive Integration of Single-Cell Data. Cell 177, 1888–1902.e21 (2019).

76. Ronneberger, O., Fischer, P. & Brox, T. U-net: Convolutional networks for biomedical image segmentation. in Lecture Notes in Computer Science (including subseries Lecture Notes in Artificial Intelligence and Lecture Notes in Bioinformatics) 9351, 234–241 (Springer Verlag, 2015).

77. Saka, S. K. et al. Immuno-SABER enables highly multiplexed and amplified protein imaging in tissues. Nat. Biotechnol. 37, 1080–1090 (2019).

78. McQuin, C. et al. CellProfiler 3.0: Next-generation image processing for biology. PLOS Biol. 16, e2005970 (2018).

